# *Toxoplasma gondii* secreted effectors co-opt host repressor complexes to inhibit necroptosis

**DOI:** 10.1101/2020.08.31.275339

**Authors:** Alex Rosenberg, L. David Sibley

**Author notes:** Correspondence: L. David Sibley.

## Abstract

During infection, *Toxoplasma gondii* translocates effector proteins directly into its host cell to subvert various signaling pathways. Here we characterize a novel secreted effector that localizes to the host cell nucleus where it modulates NCoR/SMRT repressor complex levels to repress interferon regulated genes involved in cell death. Type I and type II interferons upregulate many genes including protein kinase R (PKR), inducing formation of the necrosome complex that activates Mixed Lineage Kinase Domain Like Pseudokinase (MLKL) to execute necrotic cell death. Toxoplasma NCoR/SMRT modulator (TgNSM) acts together with another secreted effector TgIST, previously shown to down-modulate IFN-γ signaling to block immune functions. Together TgNSM and TgIST block IFN driven expression of PKR and MLKL, thus preventing host cell necroptotic death. The mechanism of action of TgNSM highlights a previously unappreciated role of NCoR/SMRT in regulation of necroptosis, assuring survival of intracellular cysts, and maintenance of chronic infection.

## Introduction

*Toxoplasma gondii* is a widespread apicomplexan intracellular parasite of warm-blooded animals that also commonly infects humans. It has been found worldwide, and nearly one third of humans are chronically infected with the parasite. Although most infections are benign, toxoplasmosis causes devastating disease in immunocompromised patients and infants infected *in-utero* (Furtado et al., 2011). Disease in immunocompromised individuals is thought to largely result from the reactivation of a chronic infection that persists through the ability of the parasite to differentiate within neuronal and muscle cells from the actively dividing lytic tachyzoite stage to an encysted, slowly growing, bradyzoite stage (Jeffers et al., 2018). During host cell invasion, the parasite establishes a replication permissive niche - the parasitophorous vacuole (PV) that is separated from the host cytoplasm by a membrane (PVM) enabling parasite growth and multiplication (Martin et al., 2007).

*T. gondii* subverts diverse host cellular functions through the secretion of multiple effectors into the host cell (Hakimi et al., 2017; Hunter and Sibley, 2012). Parasite effectors are released from two specialized sets of secretory organelles including the rhoptries and dense granules. Rhoptry bulb proteins contained within the rhoptries (i.e. ROP16, ROP17, ROP18 and ROP5) are secreted by *T. gondii* into host cell cytoplasm, typically at the onset of cell invasion before complete internalization (Hunter and Sibley, 2012). In contrast, dense granule proteins (GRAs) are secreted once parasites reside within the vacuole and then are either targeted to the PV membrane (i.e. GRA15, MAF1), the host cytoplasm (i.e. GRA18, TgWIP) or the host nucleus (i.e. GRA16, GRA18, GRA24, TgIST) (Wang et al., 2020). Successful transversion of the PVM by GRA effectors relies on proteolytic processing by aspartyl protease ASP5 and a translocon complex located on the PVM and composed of MYR1, MYR2, and MYR3 proteins (Rastogi et al., 2019).

Type II interferon-γ (IFN-γ) is required for immune control of *Toxoplasma* infection during both acute and chronic infection (Suzuki et al., 1988). Mice lacking IFN-γ receptors (Yap and Sher, 1999) are extremely susceptible to *T. gondii* infection, and a drop in IFN-γ level associates with cerebral toxoplasmosis in AIDS patients (Meira et al., 2014; Pereira-Chioccola et al., 2009). IFN-γ drives numerous antimicrobial responses including degradation of tryptophan in human cells (Pfefferkorn, 1984), induction of nitric oxide synthase (NOS2), which is primarily important in murine macrophages (Stuehr et al., 1989), upregulation of reactive oxygen species, which occurs both in human and mouse (Nathan et al., 1983). In murine cells, IFN-γ - mediated upregulation of a family of immunity related GTPases (IRGs) (Taylor et al., 2007) and guanylate binding proteins (GBPs) (Kim et al., 2012) contribute to parasite clearance, while in human cells only select GBPs have been shown to play a role (Fisch et al., 2019). We have also recently shown that type I interferon (IFN-β) is also important for the control of *T. gondii* infection in the brain during chronic infection in mice (Matta et al., 2019).

IFN-γ activates JAK1 and JAK2 to phosphorylate homodimers of STAT1 and induce expression of genes that contain a gamma-activated sequence (GAS) in their core promoters (Mostafavi et al., 2016). Similarly, IFN-β induces JAK1 and TYK2 kinases to phosphorylate STAT1/STAT2 heterodimers that associate with IRF9 to activate genes that contain a canonical IFN-sensitive response element (ISRE) in their promoters (Mostafavi et al., 2016). *T. gondii* blocks the induction interferon stimulated genes (ISGs) in response to both type I and II interferons through the secretion of a GRA protein called TgIST (Gay et al., 2016; Matta et al., 2019; Olias et al., 2016). This effector is targeted to the host cell nucleus where it blocks STAT1/STAT1 and STAT1/STAT2/IRF9 transcription through the recruitment of the Mi-2/NuRD repressive complex to the promoters of ISGs (Gay et al., 2016; Matta et al., 2019; Olias et al., 2016).

In addition to activating immune defenses, Type I and II interferons have been shown to regulate a form of regulated necrosis called necroptosis (Galluzzi et al., 2017). Necroptosis has likely evolved to counteract viral suppressors ability to block extrinsic apoptosis controlled by Caspase 8 (Kaiser et al., 2013). In the absence of active Caspase 8, interferons transcriptionally activate the RNA-responsive protein kinase PKR in a JAK1/STAT1 dependent manner. PKR activation in turn induces the formation of the necrosome complex consisting of receptor-interacting serine-threonine kinase 1 (RIPK1) and receptor-interacting serine-threonine kinase 3 (RIPK3) (Thapa et al., 2013). Together they activate the pro-necroptotic protein mixed lineage kinase domain-like (MLKL) via phosphorylation, which triggers necroptotic plasma membrane permeabilization (Galluzzi et al., 2017; Linkermann and Green, 2014). Infection by *T. gondii* has been shown to inhibit multiple caspases including Caspase 8 (Payne et al., 2003; Vutova et al., 2007). Alternatively, IFN-γ treatment of human fibroblasts infected with *T. gondii* results in their death (Niedelman et al., 2013; Seizova et al., 2019). However, the exact cell death pathways and the involvement of the JAK1/STAT signaling in these events are currently unknown.

In the present study we have utilized a spatially-restricted biotin tagging approach named APEX2 (Lam et al., 2015) to globally identify *T. gondii* effectors that are secreted into the host cell nucleus. We identified all of the formerly known nuclear effectors and a new factor that increases the level and activity of the nuclear receptor co-repressor (NCoR) and silencing mediator of retinoic acid and thyroid hormone receptors (SMRT) NCoR/SMRT repressor complexes (Mottis et al., 2013). NCoR/SMRT functions to maintain basal repression of a subset of genes that are activated by Toll-like receptors and other pro-inflammatory signaling pathways (Glass and Saijo, 2010). The parasite effector, named Toxoplasma NCoR/SMRT modulator (TgNSM) is critical during infection of the chronic bradyzoite stage, where in concert with TgIST, it blocks interferon induced necroptosis, thus prolonging cell survival and protecting the parasite’s intracellular niche.

## Results

### Utilization of the APEX2 system uncovers novel *T. gondii* secreted effectors

To find novel *T. gondii* effectors that were targeted into the host nucleus, we used the technique of spatially restricted enzymatic labeling with the engineered peroxidase APEX2 (Lam et al., 2015). This approach uses the enzyme APEX2 targeted to various subcellular locations to promote rapid (~1 min) biotin labeling of neighboring proteins within ~20nm radius (which can be subsequently purified on streptavidin and identified with mass spectrometry (MS)) (Martell et al., 2012). HeLa cells transiently expressing APEX2 either in the host cell cytoplasm or in the nucleus were infected with type I (RH) strain *T. gondii* parasites for 24 hr (Figure 1A). Untransfected HeLa cells were infected as well and used as a background control. To initiate labeling, cells were incubated with hydrogen peroxide and a biotinylated tyramide derivative (biotin-phenol). The biotinylated proteins were collected using streptavidin-coated beads and identified by MS/MS analysis. Analysis of nuclear samples from two biological experiments revealed the presence of the majority of previously identified secreted *T. gondii* effectors that traffic to the host nucleus including GRA28, GRA16, GRA24, and TgIST (Figure 1B, Table S1) (Bougdour et al., 2013; Braun et al., 2013; Gay et al., 2016; Nadipuram et al., 2016; Olias et al., 2016). We compared total unique peptide counts for the known secreted effectors identified in the cytoplasmic, nuclear, and control samples and adopted the following criteria for identifying novel effectors: 1) identified by at least two peptides in the APEX2 samples from both experiments without being detected in the control experiments, 2) demonstrated at least a 3-fold enrichment in the peptide count in nuclear versus the cytoplasmic fraction. Further, we chose to focus on proteins that also had a predicted nuclear localization domain along with a prediction of being intrinsically disordered in their three-dimensional structure, as this property has previously been recognized as a feature of secreted GRA effectors in *T. gondii* (Hakimi et al., 2017). These criteria left us with three candidate proteins: TGME49_209850, TGME49_235140 and TGME49_239010 (Figure 1B). The localization of these proteins was tested by direct tagging of the endogenous loci in *T. gondii* with a C-terminal Ty tag (Bastin et al., 1996) using the highly efficient CRISPR system (Shen et al., 2014). Two out the three tagged proteins (TGME49_235140 and TGME49_239010) indeed showed host nuclear localization (Figure 1C), while TGME49_209850 was targeted to the parasite nucleus (Figure. S1). TGME49_239010 has recently been characterized by two independent groups as a host nucleus targeted effector mediating cyclin E upregulation and modulating NF-κB signaling via interaction with E2F3 and E2F4 transcription factors (Braun et al., 2019; Panas et al., 2019). Consequently, we decided to focus on TGME49_235140, which we have named TgNSM, based on functional characterization provided below.

**Figure 1.**
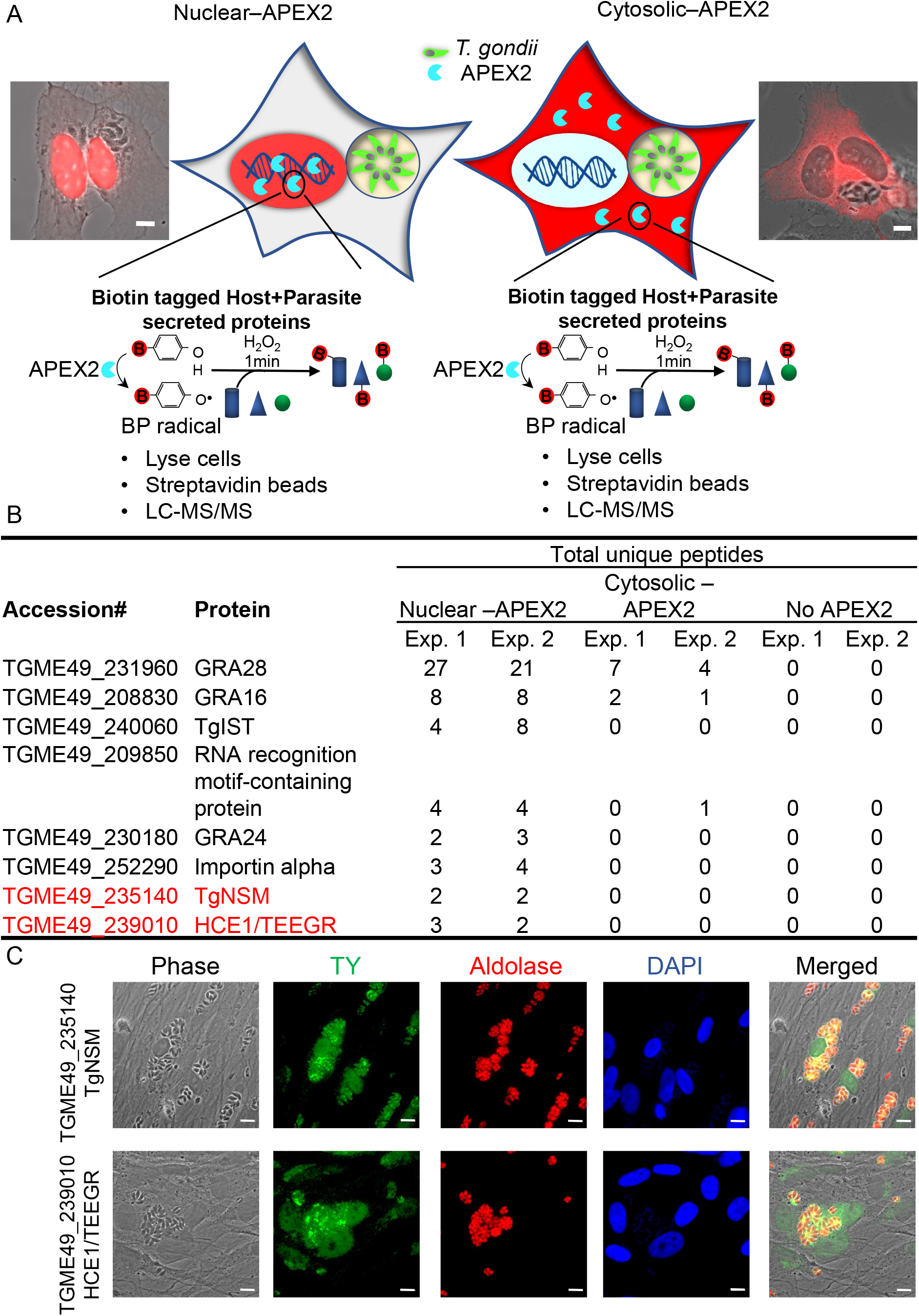
Utilizing APEX2 to uncover new effectors secreted by *T. gondii* into the host cell nucleus. (A) Schematic representation of HeLa cells expressing APEX2 in the nucleus (left) or the cytoplasm (right) and infected with *T. gondii* (green) shown in the center. Cells were treated with biotin-phenol and H_2_O_2_ to label proteins with biotin in the respective compartment (shaded red). Combined phase contrast and fluorescence images to the outside show cells stained with Alexa Fluor 568 streptavidin to detect biotin (red) labeling in the nucleus (left) and cytosol (right). Parasites are visible in phase contrast. Scale bars = 5 μm. (B) Summary of *T. gondii* proteins identified by mass spectrometry analysis of two combined APEX2 experiments with Nuclear-APEX2, Cytoplasmic-APEX2 and APEX2 free control. (C) HFF cells infected with type I (RH) endogenously tagged parasites: TGME49_235140, defined here as TgNSM-Ty, and TGME49_239010, previously described as HCE1/TEEGR–Ty (Braun et al., 2019; Panas et al., 2019). Cells were fixed 24 hr post-infection, stained with mouse anti-Ty and anti-mouse IgG Alexa Fluor 488 (green) to detect the secreted effectors, rabbit anti-Aldolase and rabbit IgG Alexa Fluor 568 (red) to detect all parasites, and DAPI (blue). Scale bars = 5 μm.

### TgNSM is a late exported dense granule protein targeted to host cell nucleus

TgNSM encodes a ~48 kDa protein that lacks a signal peptide and has a predicted single transmembrane domain near the N-terminus (Figure. 2A). TgNSM also has a potential nuclear localization signal (NLS) (Figure. 2A) and it constitutes an intrinsically disordered protein (IDP) with no domain homologies to other known proteins (Figure 2B). Immunofluorescence staining revealed that TgNSM had a punctate distribution in the parasite cytoplasm that partially colocalized with the dense granule protein GRA2 (Figure. 2C). Based on time course infection experiments, TgNSM was secreted at late time points of infection (Figure. 2C). In the first 8 hr after infection, it was mainly detected inside the parasite; however, at 20 hr post infection it was readily observed inside the host nucleus (Figure. 2C). We also infected Human foreskin fibroblasts (HFF) cells and cultured them for 5 days under alkaline conditions that induce differentiation of bradyzoites (Weiss et al., 1995). TgNSM was readily detected in host nuclei containing 5-day-old bradyzoites (Figure. 2D). Similarly to previously characterized exported GRA proteins, the translocation of TgNSM across the parasitophorous vacuole membrane was dependent on the translocon protein MYR1 (Franco et al., 2016) and, also on the aspartyl protease ASP5 (Coffey et al., 2015), although the protein does not have a consensus TEXEL motif (Figure. 2E). A similar dependence has been previously reported for dense granule protein GRA24 which lacks TEXEL motif and yet requires ASP5 for secretion into the host nucleus (Coffey et al., 2015; Curt-Varesano et al., 2016; Franco et al., 2016; Hammoudi et al., 2015). Collectively these findings indicate that TgNSM is a member of the subfamily of dense granule-resident proteins that are exported across the PVM and stably accumulate in the host cell nucleus.

**Figure 2.**
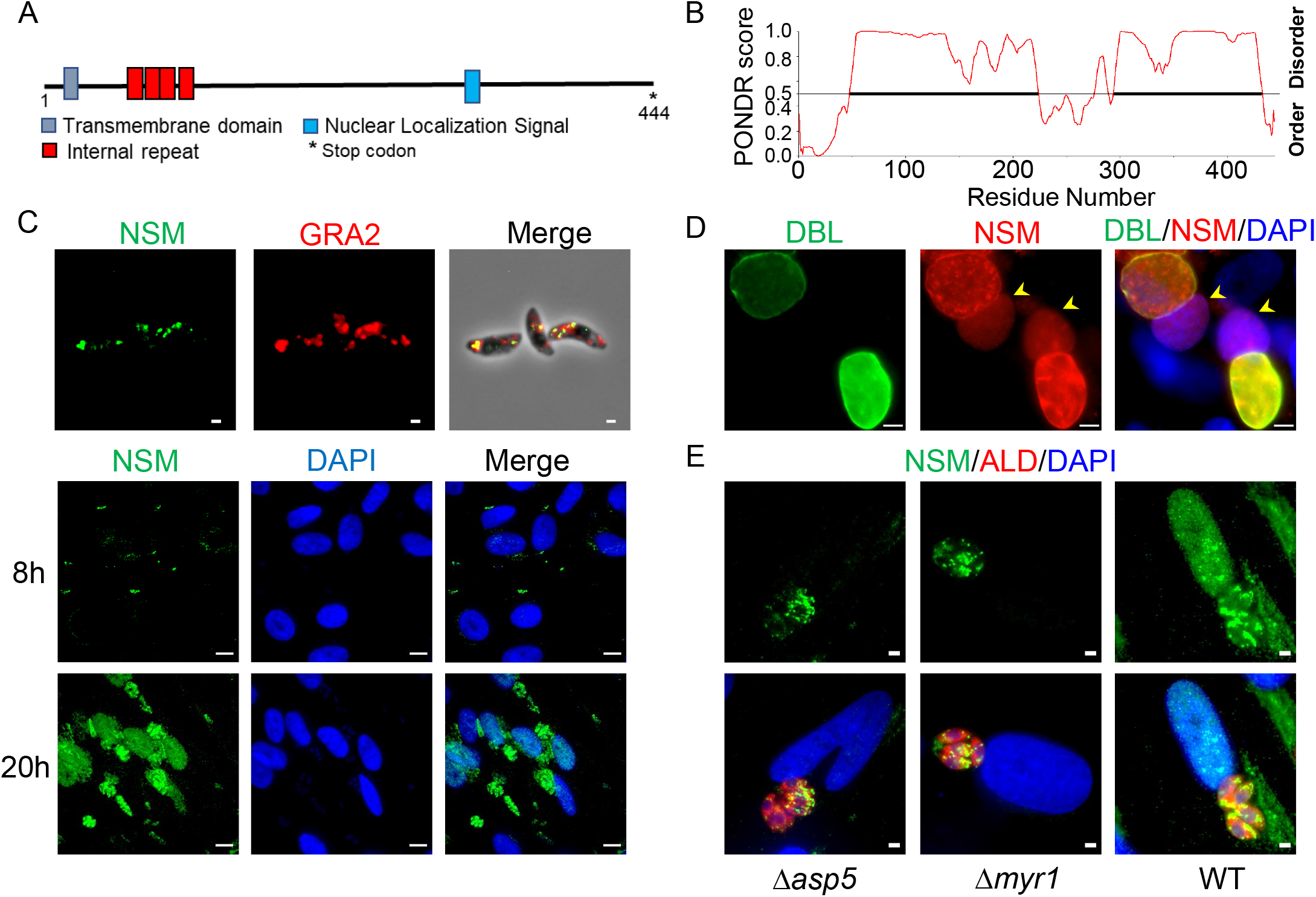
TgNSM is expressed late during infection and exported to the host cell nucleus. (A) Schematic representation of the domain structure of TgNSM including putative transmembrane domain (grey), putative nuclear localization signal (as predicted by (ELM) (cyan), and the internal repeats (red). Numbers indicate the residues from 1 to 444. (B) Schematic representation of TgNSM probability of disorder (as predicted by PONDR). Segments with values > 0.5 are predicted to be disordered, and segments with values < 0.5 correspond to folded regions. (C) Localization of TgNSM in extracellular type I (RH) tachyzoites expressing TgNSM-Ty (top panel) versus HFF cells infected with type I (RH) parasites expressing TgNSM-Ty (middle and bottom panels). Cells were fixed at intervals post-infection and stained with mouse anti-Ty and anti-mouse IgG Alexa Fluor 488 (green), rabbit anti-GRA2 and rabbit IgG Alexa Fluor 568 (red), and DAPI (blue). Scale bars = 5 μm. (D) Localization of TgNSM in HFF cells infected with ME49 Δku80 TgNSM-Ty and cultured under alkaline conditions for 5 days to induce bradyzoites before fixation. Cells were stained with biotinylated *Dolichos biflorus* lectin (DBL) and Alexa Fluor 488 Streptavidin (green), mouse anti-Ty and anti-mouse IgG Alexa Fluor 568 (red), and DAPI (blue). Arrows (yellow) point to host nuclei containing TgNSM-Ty. Scale bars = 5 μm. (E) Role of MYR1 and ASP5 in TgNSM export into the host cell. HFF cells were infected with wild type (WT) RH strain, RH *Δasp5*, or RH *Δmyr1* parasites transiently expressing TgNSM-Ty (pTgNSM-TgNSM-Ty). At 18 hr post-infection, the cultures were fixed and stained with mouse anti-Ty and anti-mouse IgG Alexa Fluor 488 (green), rabbit anti-Aldolase and rabbit IgG Alexa Fluor 568 (red), and DAPI (blue). Scale bars = 5 μm.

### TgNSM targets the host NCoR/SMRT repressor complex

To identify TgNSM targets within the host nucleus, we performed immunoprecipitation (IP) experiments utilizing a TgNSM-Ty tagged line. HFF cells were infected with either TgNSM-Ty *T. gondii* or wild type (WT) tag free parasites for 24 hr. Nuclear extracts were prepared, TgNSM-Ty was immunoprecipitated, and complexes from three independent replicates were subjected to MS analysis. Omitting all host proteins that appeared in the control fraction resulted in a list of only 12 proteins that were specifically immunoprecipitated with TgNSM-Ty (Figure 3A, Table S2). A STRING network (Szklarczyk et al., 2019) of the 12 interacting proteins contained all the core components of the NCoR/SMRT repressor complex, namely Nuclear receptor corepressor 1 (NCOR1/NCoR), Nuclear Receptor Corepressor 2 (NCOR2/SMRT), histone deacetylase 3 (HDAC3), transducin β-like 1 (TBL1), TBL-related 1 (TBLR1) and G-protein-pathway suppressor 2 (GPS2) (Figure. 3B) (Perissi et al., 2010). Further analysis of the MS data sets using the program Straightforward Filtering IndeX program (SFINX), which provides a statistically robust method for determining true interactions while filtering out false-positives from replicate data sets (Titeca et al., 2016), identified TBLR1, SMRT, TBL1 and NCOR1 as the most probable TgNSM interactors (Figure. 3C).

**Figure 3.**
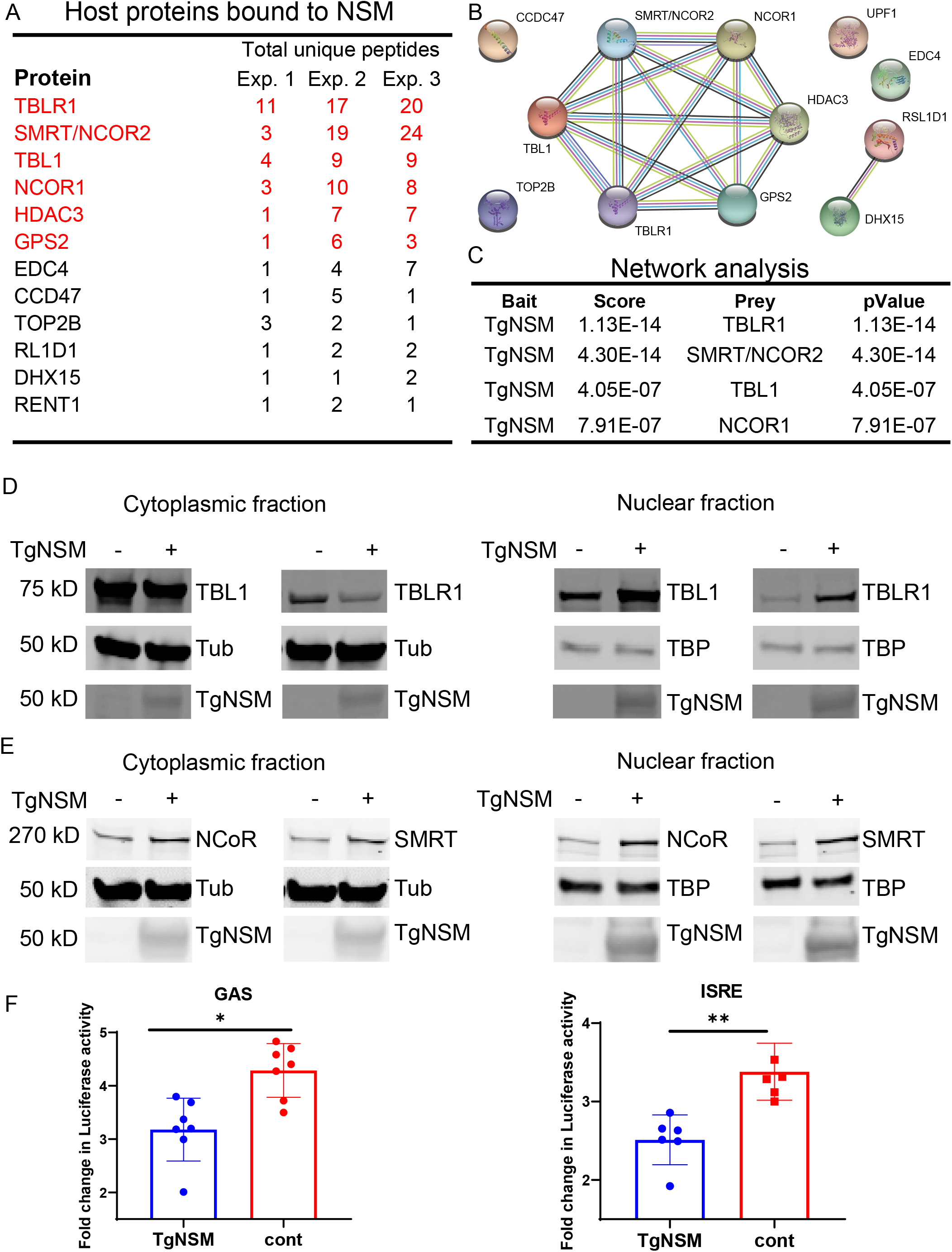
TgNSM interacts with NCoR/SMRT repressor complex components affecting their localization, expression levels, and function. (A) TgNSM-Ty-associated host proteins identified by mass spectrometry (MS) analysis. TgNSM-Ty immunoprecipitated from type I (RH) TgNSM-Ty vs. wildtype untagged type I (RH) parasite infected HFF cells. Combination of three experiments showing proteins that were solely identified in TgNSM-Ty and not in wildtype infected cells. Identity of the proteins with their respective number of peptides are indicated, NCoR/SMRT complex core components are in red. (B) Network of the TgNSM interactome using the STRING database. Proteins identified by mass spectrometry were analyzed using Scaffold version 4. 0. Proteins showing enrichment are represented by nodes (filled circles) in the network and the edges indicate interactions. Criteria for interactions are shown by colored lines: red line: gene fusion events, green line: gene neighborhood, blue line: gene co-occurrence, purple line: experimental evidence, yellow line: text mining, light blue line: database, black line: co-expression. (C) Three replicates of MS analysis data sets generated with TgNSM-Ty and WT IP experiments were analyzed using SFINX to filter out false-positive interactions and rank true-positives. TBLR1, SMRT/NCOR2, NCoR1 and TBL1 were identified as TgNSM interactors with high statistical confidence. (D) HEK 293T cells were transfected with TBL1-mCherry or TBLR1-mCherry in combination with TgNSM-Ty or empty vector. Twenty-four hr after transfection, cells were collected lysed and fractionated into cytoplasmic and nuclear fractions. The protein fractions were resolved on SDS-PAGE gels, blotted with primary antibodies, and imaged with LI-COR specific secondary antibodies. (E) HEK 293T cells were transfected with NCoR-HA or SMRT-HA in combination with TgNSM-Ty or empty vector. Twenty-four hours hr after transfection, cells were collected lysed and fractionated into cytoplasmic and nuclear fractions. The protein fractions were resolved on SDS-PAGE gels, blotted with primary antibodies, and imaged with LI-COR specific secondary antibodies. (F) GAS and ISRE Gaussia luciferase reporter constructs were transiently transfected into HeLa cells with TgNSM-Ty or empty vector. Twenty-four hours later, transfected cells were treated with IFN-γ (GAS) or IFN-β (ISRE) at 100 U/ml and 1,000 U/ml respectively for 24 hr and luciferase activity was determined. Results shown are fold induction over untreated samples and represent the averages and standard deviation from 3 biological replicates done in duplicate. Each point represents a technical replicate. Mean ± SD (n = 3 experiments, each with 2 replicates). **P < 0.01, two-sample unpaired Student’s t test.

The NCoR/SMRT complex serves as a repressive coregulatory factor (corepressor) for multiple transcription factors (Watson et al., 2012), recruiting HDAC3 to DNA promoters to regulate many developmental and metabolic pathways, as well as inflammation (Mottis et al., 2013). Based on these broad functions, we wanted to test if TgNSM influences the levels or the localization of its predicted interactors within the NCoR/SMRT complex. TBL1/TBLR1 proteins lack a canonical nuclear localization signal and are primarily cytoplasmic, but translocate to the nucleus upon stimulation (Kruusvee et al., 2017; Lyst et al., 2013; Zhang et al., 2006). Ectopic co expression of TgNSM-Ty together with TBL1 or TBLR1 in HEK 293T cells followed by subcellular fractionation resulted in translocation of TBL1 and TBLR1 into the host cell nucleus (Figure. 3D). TBL1/TBLR1 facilitate stable chromatin binding and promote the clearance of NCoR/SMRT corepressors by recruiting 19S proteasome particles to drive the ubiquitination and degradation of NCoR/SMRT, thus allowing the recruitment of coactivators (Perissi et al., 2004; Perissi et al., 2008; Yoon et al., 2003). Changes in TBLR1 levels have been shown to affect SMRT stability (Zhang et al., 2006). Therefore, we tested if TgNSM has any effect on stabilizing NCoR or SMRT proteins. We performed ectopic co-expression of TgNSM-Ty in combination with NCoR or SMRT in HEK 293T cells followed by subcellular fractionation. Analysis of the cytoplasmic and nuclear fractions revealed that the presence of TgNSM increased the levels of both NCoR and SMRT, which was more pronounced in the nuclear fraction (Figure. 3E).

The observed TgNSM mediated increase in the levels of NCoR and SMRT in the nuclear fraction prompted us to test if TgNSM can directly affect NCoR/SMRT regulated transcription of inflammatory genes. Due to importance of type I and II interferons in host response to *T. gondii*, we focused on testing the ability of TgNSM to block both IFN-γ and IFN-β responses. To this end we co-expressed luciferase reporters downstream of GAS or ISRE sequences together and without TgNSM-Ty in HeLa cells. In these experiments, the presence of TgNSM had an inhibitory effect on the transcriptional activity of both reporters upon IFN treatment (Figure 3F). Hence, TgNSM induces translocation of TBL1 and TBLR1 into the nucleus leading to increased levels of NCoR and SMRT that result in enhanced repressive activity.

### TgNSM together with TgIST inhibit IFN-γ induced necroptotic gene expression in bradyzoite infected cells

Recent studies have shown that bradyzoites lacking the MYR1 export complex (Δ*myr1* mutants), which are devoid of the ability to export any of the GRA effectors, are unable to prevent host cell death leading to cysts rupture upon IFN-γ treatment (Seizova et al., 2019). This phenotype was specific to bradyzoites and was not observed in tachyzoite infected cells, although the mechanism of cell death was not determined (Seizova et al., 2019). Surprisingly, deletion of TgIST, which is thought to be the major effector responsible for IFN-γ repression, did not phenocopy the dramatic host cell death phenotype of *Δmyr1* mutants, suggesting that an additional effector modulates similar pathway (Seizova et al., 2019). Therefore, we created single and double mutant strains in the type II (ME49) line (i.e. WT, *Δnsm, Δist and Δnsm/Δist*) (Figure S2) carrying mCardinal (mCard) driven by a constitutive tubulin promoter and the bradyzoite specific promoter BAG1 driving mNeonGreen (mNG). The strains were grown in HFF cells under alkaline conditions to induce *T. gondii* differentiation into bradyzoites for 5 days. Infected HFF monolayers were then stimulated with IFN-γ for 6 hr, harvested, and infected cells were purified by fluorescence-activated cell sorting (FACS) based on the expression of both mCard and mNG. RNA was then extracted and sequenced using Illumina chemistry on a NovaSeq platform. Biological pathway analysis of the differentially expressed genes between WT parasites and Δ*nsm*, Δ*ist*, Δ*ist*/Δ*nsm* mutants were analyzed with Reactome Pathway Knowledgebase (Jassal et al., 2020). These studies revealed that deletion of *nsm* leads to changes in p53, mTOR, and MeCP2 regulated pathways (Figure 4A). These findings are consistent with the modulation of NCoR/SMRT by TgNSM. For example, SMRT has been shown to facilitate coactivation of p53 dependent gene expression (Adikesavan et al., 2014), while NCoR activity have been shown to be affected by the mTOR signaling in the contexts of obesity and aging (Mottis et al., 2013). MeCP2 is also a known NCoR/SMRT interactor, that recruits this complex to methylated sites within the genome to downregulate transcription (Lyst and Bird, 2015). We did not detect a change in genes related to interferon signaling pathways in the Δ*nsm* mutant, likely due to the dominant repressive activity of TgIST.

**Figure 4.**
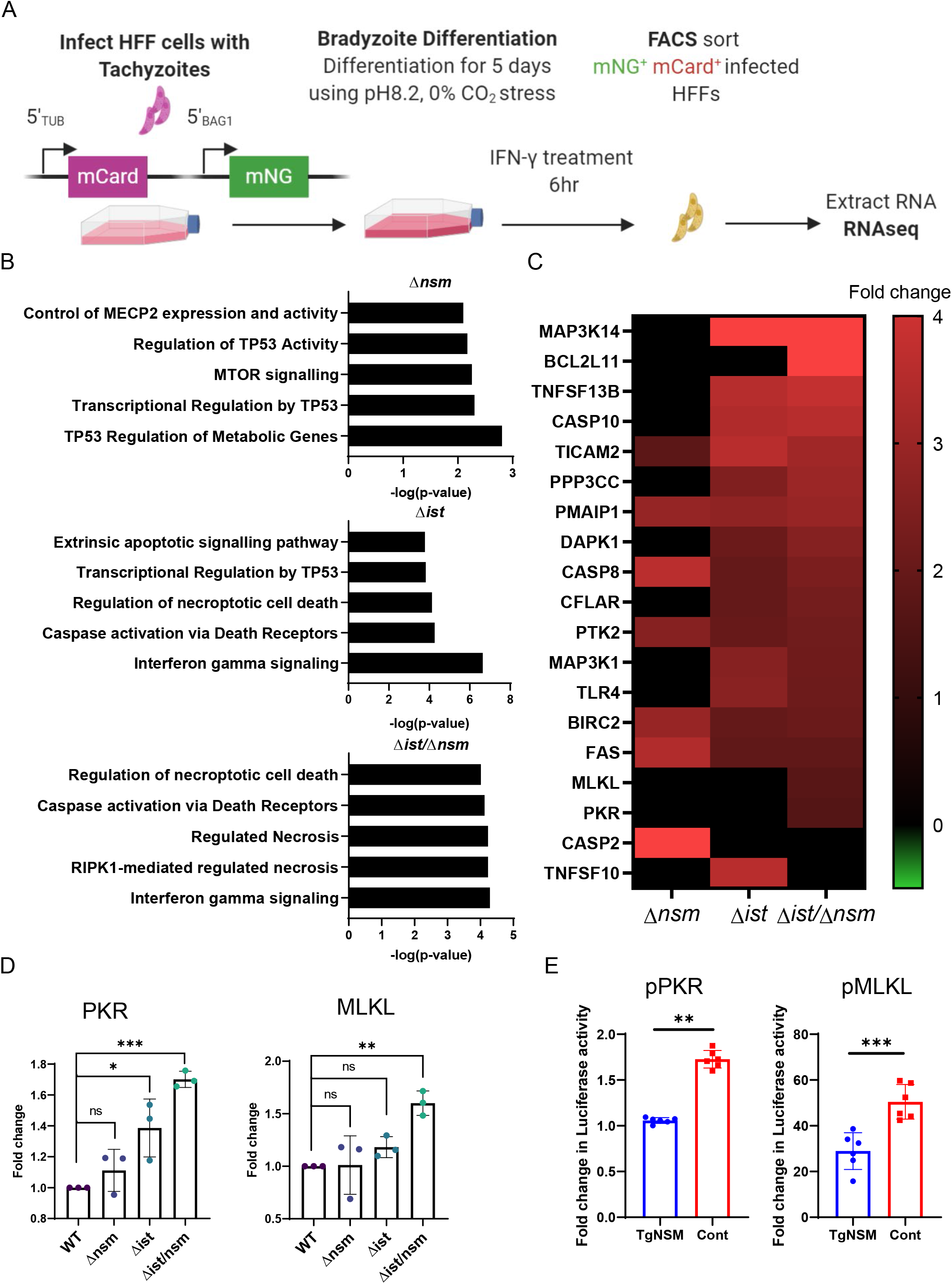
TgNSM together with TgIST block transcription of genes involved in regulated necrosis pathway. (A) Schematic of sample collection for RNA sequencing. Constitutively expressing mCardinal (mCard) cells that also express a stage-specific mNeonGreen (mNG) grown in HFF cells were induced to differentiate by culture at pH 8.2 and then sorted to obtain bradyzoites (doubly positive) infected cells. (B) Top-scoring pathways upregulated in ME49 Δ*nsm*, Δ*ist*, Δ*nsm*/*ist* mutants vs. wildtype bradyzoites upon 6 hr IFN-γ stimulation in HFFs, were determined by Reactome Pathway Knowledgebase. (C) Heat map of cell death related genes in ME49 Δ*nsm*, Δ*ist*, Δ*nsm/ist* mutants vs. wildtype. Fold change >1.5 *P* -value <0.05. (D) qPCR showing fold induction of mRNA transcripts in HFFs infected with parasite strains carrying constitutive pTUB-mCardinal and inducible pBAG1-mNeonGreen reporters, grown in HFF cells under alkaline stress for 5 days, followed by treatment with IFN-γ (100 U/mL for 6 hr) purified by FACS based on the presence of both reporters. Comparative cycle threshold (Ct) values were used to evaluate the fold change in transcripts using β-actin (ActB) as an internal transcript control. Data are plotted as fold change ± SEM from at least 3 independent experiments per gene. There were significant differences between the compared groups *\*P* <0.05, and \*\**P* < 0.01, \*\*\**P*<0.001 using one-way ANOVA with Tukey’s multiple comparison test. (E) MLKL and PKR luciferase promoter constructs were transiently transfected into HeLa cells with TgNSM-Ty or empty vector. Twenty-four hours later, transfected cells were treated with IFN-γ at (100 U/mL for 24 hr) and firefly luciferase activity was determined. The transfection efficiency was normalized against the-Rennila luciferase activity from the cotransfected pRL-TK vector. Results shown are fold induction over unstimulated control and represent the averages from 3 biological replicates done in duplicate. Each point represents a technical replicate. Mean ± SD (n = 3 experiments, each with 2 replicates). \*\*\**P* < 0.001 and ****, *P* < 0.0001, two-sample unpaired Student’s t test

When we compared Δ*ist* and Δ*ist*/Δ*nsm* mutants to WT infected cells, the number one modified pathway that differed was interferon signaling (Figure 4B). Interestingly the next top ranked pathways were related to regulated necrosis, extrinsic apoptotic signaling pathways, and caspase activation via death receptors (Figure 4B). Closer analysis of the genes ascribed to cell death pathways revealed a trend in which the Δ*ist*/Δ*nsm* showed slightly higher expression compared to the single Δ*nsm* or Δ*ist* deletion, suggestive that the parasite requires both effectors to efficiently inhibit the transcription of genes involved in cell death (Figure 4C). PKR (aka EIF2AK2) and MLKL are known direct targets for interferon induced transcription and are key components in the JAK1/STAT1 signaling cascade responsible for execution of interferon driven necroptosis upon Caspase 8 inhibition (Knuth et al., 2019; Kuhen and Samuel, 1997; Sarhan et al., 2019; Thapa et al., 2013). Given the known ability of *T. gondii* to block Caspase 8 activity (Payne et al., 2003; Vutova et al., 2007), and the critical roles of PKR and MLKL in necroptosis (Sun et al., 2012; Thapa et al., 2013) we decided to further focus on these two genes. We first validated the mRNA expression differences between the strains with real time PCR (Figure 4 D). The highest expression of both genes was observed in the Δ*nsm*/Δ*ist* strain conforming the RNAseq results. PKR and MLKL are known to be STAT1 targets, therefore we wanted to test if TgNSM alone can inhibit the expression of both these genes. To this end we co-expressed luciferase reporters downstream of PKR and MLKL promoter sequences together and without TgNSM-Ty in HeLa cells. Co-expression of TgNSM significantly inhibited the activity of the reporters upon IFN-γ or IFN-β treatment (Figure 4D, Figure S3).

### Toxoplasma blocks interferon induced necroptosis by a combinatory function of TgNSM and TgIST

To assess the effect of TgNSM and TgIST on host cell survival, parasites were grown in HFF cells under alkaline conditions for 5 days to induce bradyzoite formation. The infected HFF monolayers were then stimulated ± IFN-γ for 15 hr in the presence of Propidium Iodide (PI). Cells were then fixed and stained with *Dolichos biflorus* lectin (DBL). Host cell death upon IFN-γ treatment was first detected by PI uptake and staining of the host nucleus that was followed by cyst rupture (Figure 5A). Therefore, to quantify the effect of IFN-γ on host cell survival and cyst rupture, the number of intact (DBL positive) cysts residing within PI negative host cells was quantified by microscopy imaging and normalized to the number of intact cysts in the control monolayers not treated with IFN-γ. No significant difference in cyst survival between the WT and the single Δ*nsm* or Δ*ist* mutants was observed (Figure 5B). However, double Δ*nsm*/Δ*ist* mutants exhibited a large drop in cyst survival that was significantly different from the WT and single mutants (Figure 5B). The host cell death and cyst rupture phenotype was rescued by complementing Δ*nsm*/Δ*ist* mutant with either *NSM* or *IST* (Figure 5B). Similar results were observed when the cyst infected cells were treated with IFN-β (Figure S4A).

**Figure 5.**
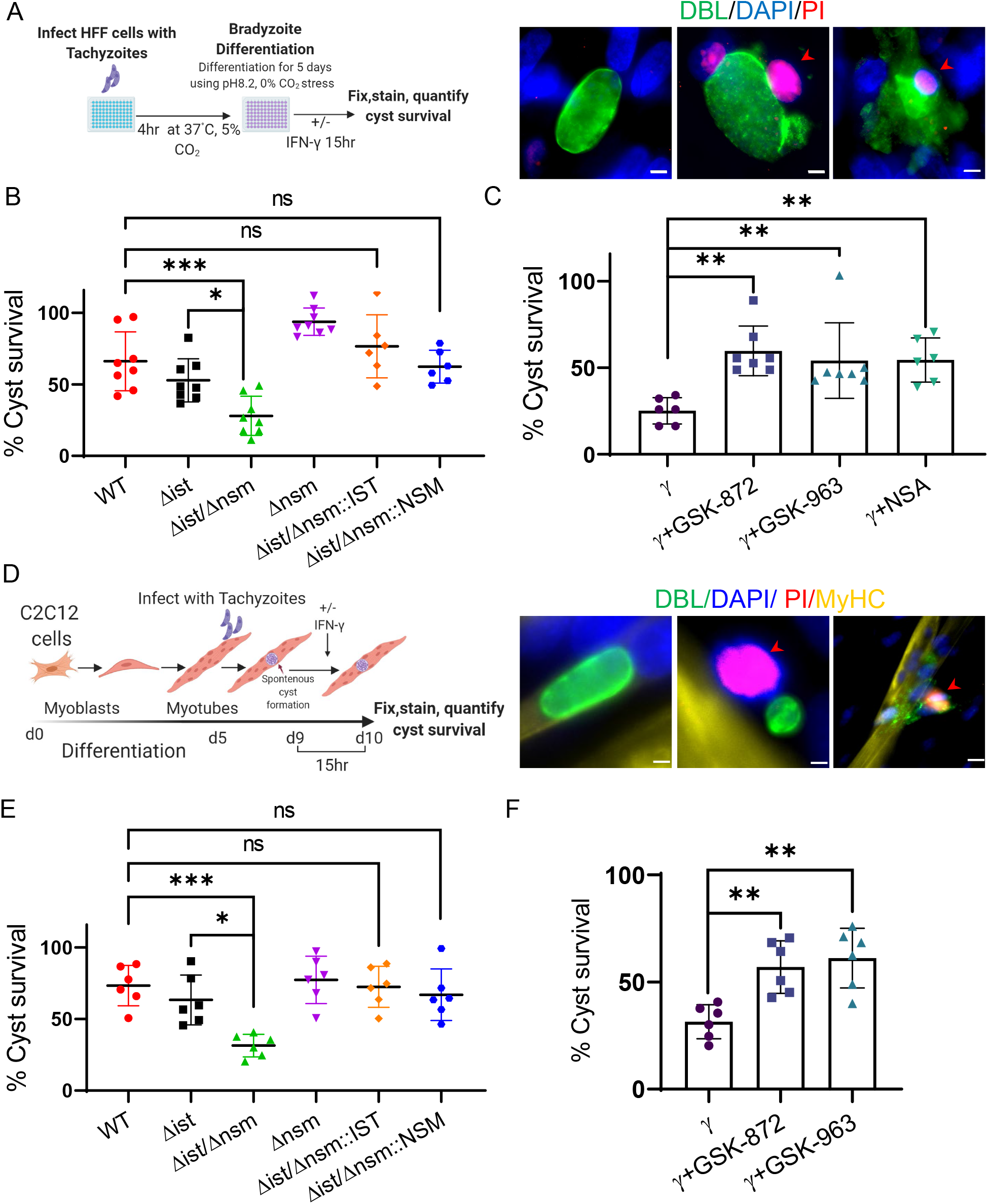
Deletion of TgNSM and TgIST induces necroptotic host cell death and cyst rupture upon IFNγ stimulation. (A) Schematic illustration of the workflow used to examine bradyzoite viability. Combined representative fluorescence images used to generate the data shown in panels B and C. HFF cells infected with ME49 Δ*nsm*/Δ*ist* and kept under alkaline conditions for 5 days to induce differentiation. Infected monolayers were treated with ± IFN-γ 100 U/ml for 15 hr in the presence of Propidium Iodide (PI 50 μg/ml) before fixation and IF staining. Intact cyst (left), intact cyst within PI positive host cell (middle), ruptured cyst (right), biotinylated *Dolichos biflorus* lectin (DBL) and Alexa Fluor 488 Streptavidin (green), PI (red) and DAPI (blue). Arrows (red) point to PI positive host nuclei. Scale bars = 5 μm. (B) Parasites were grown and treated as in A and the survival of cysts was calculated as percent of intact/PI negative cysts in IFN-γ treated samples compared to IFN-γ free control and represent the averages and standard deviation from at least 3 biological replicates with each dot representing a technical duplicate, Mean ± SD (n = 3 experiments, each with 2 replicates 50-100 parasite cysts were counted in each treatment). *\*P* < 0.05, *** *P* < 0.001, one-way ANOVA test with Tukey’s multiple comparison test. (C) ME49 Δ*nsm*/Δ*ist* parasites were differentiated under alkaline conditions for 5 days before IFN-γ 100 U/ml treatment in presence PI 50 μg/ml and RIPK1 inhibitor GSK’963 1 μM, or RIPK3 inhibitor GSK’872 5 μM, or MLKL inhibitor NSA 10 μM. PI and DMSO 0.05% were used in control wells. Results shown are survival of cysts upon IFN-γ treatment, calculated as percent of intact/PI negative cysts, compared to IFN-γ free control and represent the averages and standard deviation from at least 3 biological replicates with each dot representing a technical duplicate Mean ± SD (n = 3 experiments, each with 2 replicates 50-100 parasite cysts were counted in each treatment). \*\**P* < 0.01, one-way ANOVA test with Dunnett’s multiple comparison test. (D) Schematic illustration of the workflow used to examine bradyzoite viability in muscle cells. Combined representative fluorescence images used to generate the data shown in panels E and F. C2C12 cells were differentiated to mature myotubes for 5 days and then infected with ME49 Δ*nsm*/Δ*ist* for 4 days prior to ± IFN-γ 100 U/ml treatment for 15 hr in the presence of Propidium Iodide (PI 50 μg/ml) before fixation and IF staining. Intact cyst (left), intact cyst within PI positive host cell (middle), ruptured cyst (right), Biotinylated *Dolichos biflorus* lectin (DBL) and Alexa Fluor 488 Streptavidin (green), PI (red), mouse anti myosin heavy chain MyHC anti-mouse Alexa Fluor 647 (yellow) and DAPI (blue). Arrows (red) point to PI positive host nuclei, scale bars = 5 μm. (E) C2C12 cells differentiated into myotubes were infected with parasites according to the scheme shown in D. Survival of cysts upon IFN-γ treatment was calculated as percent of intact/PI negative cysts compared to IFN-γ free control and represent the averages and standard deviation from at least 3 biological replicates with each dot representing a technical duplicate, Mean ± SD (n = 3 experiments, each with 2 replicates 50-100 parasite cysts were counted in each treatment). \**P*<0.05, *** *P* < 0.001, one-way ANOVA test with Tukey’s multiple comparison test. (F) C2C12 cells differentiated into myotubes were infected with ME49 Δ*nsm*/Δ*ist* parasites for 4 days before IFN-γ 100 U/ml treatment in presence of PI 50 μg/ml and GSK’963 1 μM or GSK’872 5 μM of PI 50 μg/ml. PI and DMSO 0.05% were used in control wells. Survival of cysts upon IFN-γ treatment calculated as percent of intact/PI negative cysts compared to IFN-γ free control and represent the averages and standard deviation from at least 3 biological replicates with each dot representing a technical duplicate Mean ± SD (n = 3 experiments, each with 2 replicates 50-100 parasite cysts were counted in each treatment). \*\**P* < 0.01, one-way ANOVA test with Dunnett’s multiple comparison test.

Necroptosis is driven by activation of RIPK1 followed by a formation of RIPK1-RIPK3 complex leading to activation of MLKL and subsequent cell death. Chemical inhibition of each one of these components has been shown to block the necroptotic cell death (Nogusa et al., 2016; Sun et al., 2012). Motivated by our transcriptional data, we next examined the survival of Δ*nsm*/Δ*ist* cysts after IFN-γ treatment in presence of RIPK1, RIPK3 and MLKL specific inhibitors GSK’963, GSK’872 and Necrosulfonamide (NSA), respectively. Addition of each one of the inhibitors to IFN-γ treated monolayers inhibited the loss of Δ*nsm*/Δ*ist* cysts observed in the absence of inhibitors (Figure 5C). Addition of RIPK1, RIPK3 and MLKL inhibitors also rescued host cell death and cyst rupture upon IFN-β treatment (Figure S4B). We also tested inhibitors of other cells death pathways (Apoptosis-Z-VAD, Ferroptosis-Ferrostatin-1 (Fer-1), Autophagy-3-Methyladenine (3-MA) and Pyroptosis – Z-YVAD). Addition of any of these inhibitors concomitant with IFN-γ did not improve survival of host cells or cysts after IFN-γ treatment (Figure S4C), supporting the conclusion that death pathway observed here was indeed necroptosis.

Fibroblasts are not the natural niche for differentiation of *T. gondii* cysts, and alkaline stress used to induce bradyzoite differentiation might affect host cell natural response to the parasite. Therefore, we wanted to test if the TgNSM and TgIST also function to block cell death in the natural setting of chronic infection. *T. gondii* forms cyst spontaneously upon infection of neuronal and muscle cells. We examined the fate of cysts in C2C12 myoblasts derived from murine skeletal muscle cells that can be differentiated into myotube cells and support spontaneous differentiation into bradyzoites (Swierzy et al., 2014; Takacs et al., 2012). C2C12 myoblasts were differentiated for 5 days prior to infection, 4 days after infection the cells were stimulated ± IFN-γ for 15 hr in the presence of Propidium Iodide (PI). At the end of 15 hr period the cells were fixed and stained with DBL and myosin heavy chain (MyHC) that is indicative for terminal differentiation of C2C12 myoblasts. The number of intact/PI negative cysts was quantified using microscopy imaging (Figure 5D). The double Δ*nsm*/Δ*ist* mutant exhibited a significant drop in host cell survival and intact cyst while the single Δ*nsm* and Δ*ist* mutants were not significantly different from the WT (Figure 5E). The cell death phenotype was rescued by complementation with either single *NSM* or *IST* (Figure 5E). Inhibition of RIPK1 and RIPK3 reversed the sensitivity to interferon induced cell death and cysts rupture (Figure 5F). Collectively, these results indicate that *T. gondii* bradyzoites rely on a combinatorial inhibitory function of TgNSM and TgIST to prevent interferon induced necroptotic host cell death.

## Discussion

*Toxoplasma* modulates host gene expression to alter a number of pathways, including the ability to prevent cell death, although the mechanism of this blockade is unknown. Here we utilized the APEX2 system to identify a novel *T. gondii* secreted effector TgNSM that localizes to the host cell nucleus and functions to block the necrotic death pathway. TgNSM drives nuclear translocation of TBL1/TBLR1 that function as co-repressor-co-activator exchange factors leading to an increase in both NCoR and SMRT nuclear levels and their subsequent transcriptional repression of interferon stimulated genes. In addition to their role in immunity, type I (α/β) and type II (γ) IFNs induce necroptosis through Jak1/STAT1-driven transcription of PKR that induces the formation of the necrosome complex leading to necrotic cell death that is mediated by MLKL. We show that TgNSM functions in concert with TgIST to inhibit IFN driven expression of PKR and MLKL, thus blocking cyst infected host cell necroptotic death. The ability of *T. gondii* to prevent necrotic death is important for endurance of tissue cysts that mediate chronic infection, thus ensuring parasite survival and transmission.

Previously, identifying Toxoplasma effectors exported from the parasite in vivo was limited by the fact that they constitute only a small fraction of the host cell proteome. Here we used the sensitivity of APEX2 expression within the host cell to limit the background labeling of parasite proteins, allowing us to comprehensively identify secreted effectors. We detected all the known effectors that traffic to the nucleus as well as several novel ones. Although here we focused on nuclear targeted effectors, *T. gondii* also secretes effectors to other cellular compartments. For example GRA18 and TgWIP are secreted into the cytosol (He et al., 2018; Sangare et al., 2019), while MAF1 directly interacts with host mitochondria (Pernas et al., 2014), and it is possible that other organelles represent additional destinations for the secreted effectors. Hence, approaches such as APEX2 could be applied towards discovery of additional effectors targeted to various organelles and cytoplasm. APEX2 could also be implemented for discovery of effectors secreted during other life cycles stages, for example bradyzoite stages. Further, this experimental approach would be beneficial for studying secreted effectors within other genetically intractable pathogenic organisms.

Similar to previously characterized secreted effectors that traffic to the host nucleus (Hakimi et al., 2017; Rastogi et al., 2019), successful delivery and translocation of TgNSM across the PVM depends on aspartic protease TgASP5 and the translocon machinery component TgMYR1 (Wang et al., 2020). TgNSM is secreted late and could also be detected in host nuclei 5 days post infection under bradyzoite growth conditions. Although we cannot be sure if TgNSM was initially secreted by tachyzoites and simply remained stable in the host nucleus *vs*. being continuously secreted, it is comparable to TgIST secretion dynamics under the same conditions (Mayoral et al., 2020; Seizova et al., 2019).

*T. gondii* uses TgNSM together with TgIST as fail-safe mechanism to block expression of a set of genes involved in regulated necrosis. Within this set of genes, we found two key components of IFN driven necroptosis - PKR and MLKL. Using ectopic expression and gene depletion experiments we reveal that TgNSM is both sufficient and necessary for the efficient repression of PKR and MLKL by bradyzoites. TgNSM and TgIST were also critical in maintaining viability of differentiated muscle cells in presence of IFN-γ, hence this pathway is also relevant in cell types where tissue cysts normally develop and persist in vivo. Differentiation to the cyst form is an adaptation to remain within the host cells for prolonged periods in order to escape immunity and assure transmission to subsequent hosts (Watts et al., 2015). Therefore, maintaining host cell viability is especially critical for the chronic form of the parasite.

Our studies demonstrate that chemical inhibition of the necroptotic pathway (RIPK1, RIPK3 and MLKL), but not other cell death pathways, prevented cell death and rupture of intracellular cysts. RIPK3 can also activate caspase-1-dependent pyroptosis-a highly inflammatory form of programmed cell death that has been ascribed to the resistance of the Lewis rat to *Toxoplasma* (Cirelli et al., 2014; Gorfu et al., 2014). However, chemical inhibition of Caspase-1 did not rescue the death phenotype in the Δ*nsm*/Δ*ist* mutant indicating that pyroptotic death pathway is not important for cell death and rupture of tissue cysts under conditions tested here. Although our studies demonstrate a role for necrotic cell death under conditions where tissue cysts form under alkaline conditions and during natural differentiation in muscle cells, the significance of this phenotype in vivo is still uncertain. During acute infection, ripk3^-/-^ and mlkl^-/-^ mice are equally or even less sensitive to infection with type II strains of *T. gondii* (Cervantes and Knoll, 2020; DeLaney et al., 2019). However, our studies predict that the main role for TgNSM in modulating the necroptotic cell death pathway is likely during chronic infection in vivo when the parasite persists in tissue cysts in muscles and neurons. Although we have not been able to test this hypothesis directly, owing to the severe attenuation of the Δ*ist* mutant (Gay et al., 2016), the continued expression of TgNSM (present report) and TgIST (Mayoral et al., 2020) within differentiated bradyzoites is consistent with a role in chronic infection.

TgIST drives the recruitment of Mi-2/NuRD to interact with the phosphorylated STAT1 dimers on transcriptional start sites that normally respond to INF-γ thus silencing transcription (Gay et al., 2016; Olias et al., 2016). This interaction is unique to infection and there is normally no interaction between these two complexes in non-infected cells. In contrast, TgNSM capitalizes on the preexisting ability of NCoR/SMRT to inhibit inflammatory genes (Ghisletti et al., 2009) by further enhancing its repressive activity. Within the context of cell death, SMRT has been shown to repress a set of pro-apoptotic genes, protecting from genotoxic stress-induced caspase activation (Scafoglio et al., 2013). Furthermore, the NCoR/SMRT complex was shown to be a target of caspase proteolysis during apoptosis (Mahrus et al., 2008). Thus, our results show a potential unrecognized role for the NCoR/SMRT in necroptotic death regulation. Our study demonstrates that parasites manipulate host transcription not only to block induction of immunity but to modulate cell death pathways that favor survival and thus assure long term persistence and transmission. Unlocking the mechanism behind these events could thus be used to fine tune immunity and cell death survival pathways, not only to combat pathogens but also to augment human health.

## Acknowledgements

We thank Jennifer Barks for technical assistance with tissue culture. This work was partially supported in part by grants from the National Institutes of Health to L.D.S. (AI118426). A.R. was partially supported by a Berg Postdoctoral Fellowship from the Department of Molecular Microbiology at Washington University in St. Louis. We thank Alice Ting, Martin Privalsky, Atlanta Cook, Yong-Soo Bae, Stefan Wirtz, and Conrad C. Weihl for generously providing reagents and members of the Sibley laboratory for helpful suggestions. Proteomic studies were performed by Dr. Michael Naldrett at the Proteomics and Metabolomics Facility, Center for Biotechnology at the University of Nebraska-Lincoln.

## Author Contributions

Conceptualization, A.R and L.D.S.; Methodology, A.R. and L.D.S.; Investigation, A.R; Formal Analysis, A.R; Writing – Original Draft, A.R and L.D.S.; Writing – Review & Editing, L.D.S. and A.R; Funding Acquisition, L.D.S.; Resources, L.D.S.; Supervision, L.D.S.

## STAR Methods

## RESOURCE AVAILABILITY

### Lead Contact

Further information and requests for resources and reagents should be directed to the lead contact, L. David Sibley.

### Materials Availability

All unique/stable reagents generated in this study are available from the Lead Contact with a completed Materials Transfer Agreement.

### Data and Code Availability

The datasets generated during this study are available at Gene Expression Omnibus [GEO: GSE157018].

## EXPERIMENTAL MODEL AND SUBJECT DETAILS

### Parasite and Host Cell Culture

*T. gondii* tachyzoites were serially passaged in human foreskin fibroblast (HFF) monolayers cultured in D10 medium [Dulbecco’s modified Eagle’s medium, DMEM (Invitrogen)] supplemented with 10% HyClone fetal bovine serum (GE Healthcare Life Sciences), 10 μg/mL gentamicin (Thermo Fisher Scientific), 10 mM glutamine (Thermo Fisher Scientific)]. HeLa cells (ATCC CCL-2), HEK 293T cells (CRL-11268), were maintained in D10. C2C12 mouse myoblasts (ATCC CRL-1772) were maintained in D20 (DMEM supplemented with 20% FBS) complete media at 37°C in 5% CO_2_. All strains and host cell lines were determined to be mycoplasma negative with the e-Myco plus kit (Intron Biotechnology). Strains used in this study are listed in Table S3. Gene disruptants and complemented lines were generated using CRISPR/Cas9 (Shen et al., 2014), as described in the Method Details. Parasite lines generated in other studies include RHΔhxgprtΔku80 (Huynh and Carruthers, 2009) and ME49 FLuc (Tobin and Knoll, 2012).

## METHOD DETAILS

### Plasmid Construction and Genome Editing

Plasmids were generated by site-directed mutagenesis of existing plasmids or assembled from DNA fragments by the Gibson method (Gibson, Young et al. 2009). All plasmids used in this study are listed in Table S4.

### Primers

All primers were synthesized by Integrated DNA Technologies. All CRISPR/Cas9 plasmids used in this study were derived from the single-guide RNA (sgRNA) plasmid pSAG1:CAS9-GFP, U6:sgUPRT (Shen, Brown et al. 2014) by Q5 site-directed mutagenesis (New England Biolabs) to alter the 20-nt sgRNA sequence, as described previously (Long, Brown et al. 2017). Primers for plasmids are listed in, Table S5.

### Parasite Transfection

Following natural egress, freshly harvested parasites were transfected with plasmids, using protocols previously described (Shen et al., 2014). In brief, ~2 × 10^7^ extracellular parasites were resuspended in 370 μL cytomix buffer were mixed with ~30 μL purified plasmid or amplicon DNA in a 4-mm gap BTX cuvette and electroporated using a BTX ECM 830 electroporator (Harvard Apparatus) using the following parameters: 1,700 V, 176-μs pulse length, 2 pulses, 100-ms interval between pulses. Transgenic parasites were isolated by outgrowth under selection with mycophenolic acid (25 μg/mL) and xanthine (50 μg/mL) (MPA/Xa), pyrimethamine (Pyr) (3 mM), chloramphenicol (20 mM), 5-fluorodeoxyuracil (10 μM) (Sigma), as needed. Stable clones were isolated by limiting dilution on HFF monolayers grown in 96-well plates.

### In Vitro Bradyzoite Assay

HFF monolayers grown in 96 well glass bottom plates were infected with 2.5 x 10^3^ parasites per well in D10 medium for 2 hr at 37° C to allow for invasion. For high pH bradyzoite induction, the media was changed to 5% HyClone fetal bovine serum (GE Healthcare Life Sciences) RPMI-HEPES (Thermo Fisher) lacking sodium bicarbonate, pH 8.2, then grown for an additional 5 days at 37°C at ambient CO_2_ levels (~0.03% CO_2_). The media was changed every 2 days (to prevent acidification). Bradyzoites were either stimulated with human IFN-γ (R&D systems) at final experimental concentration of 100 U/ml or IFN-β at final experimental concentration of 1000 U/ml for 15 hr after 5 days in differentiation media or left unstimulated. Propidium Iodide (PI) (1 mg/ml; Thermo Fisher) was added to the medium at a final concentration of 50 μg/ml. For cell death inhibitors testing the 5 day old bradyzoites cultures were preincubated for 30 min with relevant inhibitor or with vehicle alone (DMSO) prior to IFN-γ/IFN-β stimulation for 15 hr. The inhibitors were also present during the15 hr IFN-γ/IFN-β stimulation.

Monolayers were washed with PBS and then fixed with 4% formaldehyde and permeabilized with 0.2% Triton X-100. The parasite vacuoles were labeled with rabbit anti-TgAldolase and counterstained with anti-rabbit IgG Alexa Fluor 568. Cysts were labeled with Biotin-conjugated *Dolichos biflorus* lectin (Vector laboratories) followed by counterstain with Alexa Fluor 488 Streptavidin. The wells were manually quantified using a Zeiss Axio Observer microscope (20x objective). Cyst survival was determined by calculating the frequency of intact DBA^+^/PI^-^ parasite vacuoles in IFN-γ or IFN-β treated vs. untreated. Each parasite line was compared to WT for statistical significance using 1-way ANOVA from at least N = 3 similar experiments with n = 2 replicates per sample that included counting at least 50-100 vacuoles per replicate.

For spontaneous tachyzoite-to-bradyzoite conversion, C2C12 myoblasts were propagated in DMEM supplemented with 20% FCS 10 μg/mL gentamicin (Thermo Fisher Scientific). Cells were seeded at 4×10^3^ per well in 96-well tissue culture plates. After 24 hr, culture medium was exchanged to DMEM, 2% horse serum (Remel) and antibiotics as above and cells were allowed to differentiate to myotubes during the next 120 hr. The differentiated myotube wells were infected with 2 x 10^3^ parasites per well in DMEM, 2% horse serum. Four days after infection the bradyzoites were stimulated with mouse IFN-γ (R&D systems) at final experimental concentration of 100 U/ml for 15 hr or left unstimulated. Propidium Iodide (1 mg/ml; Thermo Fisher) was added to the media at a final concentration of 50 μg/ml. For cell death inhibitors testing the 4 day old bradyzoites culture were preincubated for 30 min with relevant inhibitor or with vehicle alone (DMSO) prior to IFN-γ stimulation with 100 U/ml for 15 hr. The inhibitors were also present during the15hr IFN-γ stimulation. Formaldehyde fixation immune staining and cyst quantification was done similarly to HFF monolayers.

### Biotin-Phenol Labeling in Live Cells

Biotin-phenol labelling were performed as previously described (Hung et al., 2016). HeLa cells transfected using polyethylenimine (PEI) to transiently express cytosolic APEX2-NES (Lam et al., 2015) or nuclear APEX2-H2B (Lee et al., 2016) were infected with freshly egressed RHΔhxgprtΔku80. Untransfected HeLa cells were infected as well and used as a background control. Twenty four hr after infection Biotin-phenol was added to the infected monolayers at final concentration of 0.5 mM, and cells were incubated at 37°C for 30 min. Biotinylation was initiated by the addition of H_2_O_2_ to a final concentration of 1 mM for 1 min, and halted by removing the media and five consecutive washes with quencher solution (10 mM sodium azide, 10 mM sodium ascorbate, and 5 mM Trolox in phosphate-buffered saline (PBS)). HeLa cells were scraped pelleted by centrifugation for 10 min at 3,000*g* at 4 °C. Cell pellets were lysed by gentle pipetting in RIPA lysis buffer supplemented with 1× Halt™ Protease Inhibitor Cocktail (100X) (Thermo), 1 mM PMSF and quenchers (10 mM sodium azide, 10 mM sodium ascorbate and 5 mM Trolox). For enrichment of biotinylated proteins with streptavidin beads, 8 mg of protein from each pool was diluted with RIPA buffer to reach 1.8 ml total volume. Streptavidin-conjugated magnetic beads (Thermo Scientific) (223 μl of per pool) were dispensed in 2 ml microcentrifuge tubes. A magnabind magnet (Thermo Scientific) was used to separate beads from the buffer solution. Beads were washed three times in RIPA buffer, and incubated with the corresponding lysate pools over night at 4 °C with gentle rotation. The beads were then washed twice with 0.2% SDS, and then washed once with buffer 1 (20 mM Tris pH7.5 and 2% SDS), followed by washing twice with buffer 2 (0.1% DOC, 1% Triton X-100, 500 mM NaCl, 1mM EDTA and 50 mM HEPES, pH7.5), once with buffer 3 (250 mM LiCl, 0.5% NP-40, 0.5% DOC, 1 mM EDTA and 10 mM Tris pH8.1), twice with buffer 4 (50 mM Tris, pH7.4 and 50 mM NaCl) and twice with PBS. The experiment was performed as two independent biological replicates (Table S1).

### IP

HFF cells were either left uninfected or were infected for 24 hr with the parasite strain RH-TgNSM-Ty. Nuclear extracts were prepared using the NE-PER Nuclear and Cytoplasmic Extraction Reagent kit (ThermoFisher) on ice at 4°C. Anti-Ty mAb BB2 beads bound to Protein G Dynabeads (Life Technologies) were precleared and incubated with cell lysates. Beads were washed and used further for MS/MS analysis. Three independent replicates were performed (Table S2).

### MS/MS analysis

The Streptavidin Magnetic Beads (ThermoFisher) for APEX2 experiments or Protein G Dynabeads (ThermoFisher) for immunoprecipitation experiments were placed in ammonium bicarbonate, then reduced (2 mM DTT for 1 hr at 37°C) and alkylated (10mM IAM for 20 min at 22°C in the dark). One μg of sequencing grade trypsin (Promega) was then added per sample and digestion was carried out overnight at 37°C. Of the final 120 μL digest, 60 μL was dried down and redissolved in 30 μL of 2.5% acetonitrile, 0.1% formic acid. Five μL of each digest was run by nanoLC-MS/MS using a 2 hr gradient on a 0.075mm x 250mm C18 Waters CSH column feeding into a Q-Exactive HF mass spectrometer.

All MS/MS samples were analyzed using Mascot (Matrix Science, London, UK; version 2.5.1). Mascot was set up to search SwissProt 2016_11, Homo sapiens (20,130 entries), the ToxoDB-28_TgondiiME49_Annotated Proteins 20160816 (8322 entries) and the cRAP_20150130 database (117 entries) assuming the digestion enzyme trypsin. Mascot was searched with a fragment ion mass tolerance of 0.060 Da and a parent ion tolerance of 10.0 PPM. Carbamidomethyl of cysteine was specified in Mascot as a fixed modification. Deamidated of asparagine and glutamine and oxidation of methionine were specified in Mascot as variable modifications. Scaffold (version Scaffold_4.7.5, Proteome Software Inc., Portland, OR) was used to validate MS/MS based peptide and protein identifications. Peptide identifications were accepted if they could be established at greater than 80.0% probability by the Peptide Prophet algorithm (Keller et al., 2002) with Scaffold delta-mass correction. Protein identifications were accepted if they could be established at greater than 99.0% probability and contained at least 2 identified peptides. Protein probabilities were assigned by the Protein Prophet algorithm (Nesvizhskii et al., 2003). Proteins that contained similar peptides and could not be differentiated based on MS/MS analysis alone were grouped to satisfy the principles of parsimony. Proteins sharing significant peptide evidence were grouped into clusters.

### Mass spectrometry and interactome analysis

The data sets in Scaffold (v4.7.5) were filtered with min #peptide = 2, protein threshold ≤99%, and peptide threshold ≤95%. The resulting hit lists were exported to excel and edited to match the format with the SFINX analysis tool (http://sfinx.ugent.be) (Table S2). A separate excel file containing NSM bait was uploaded to software SFINX website (http://sfinx.ugent.be/) together with the mass spectrometry data file.

### Western Blotting

HEK293T were transiently transfected with either SMRT-HA, NCoR-HA, TBL1-mCherry, TBLR1-mCherry in combination with empty vector or NSM-TY (Table S4) using Mirus - TransIT-LT1 Transfection Reagent. Cytoplasmic and nuclear extracts were prepared using the NE-PER Nuclear and Cytoplasmic Extraction Reagent kit (ThermoFisher) on ice at 4°C. Protein samples were prepared in Laemmli buffer containing 100 mM dithiothreitol, boiled for 5 min, separated on 8%–16% polyacrylamide gels by SDS-PAGE, and transferred to nitrocellulose membranes. The membranes were blocked with 5% (wt/vol) fat-free milk in PBS and then probed with primary antibodies diluted in blocking buffer containing 0.1% Tween 20. Membranes were washed with PBS + 0.1% Tween 20, then incubated with goat IR dye-conjugated secondary antibodies (LI-COR Biosciences) in blocking buffer as indicated in the figure legends or associated method details. Membranes were washed several times before scanning on a Li-Cor Odyssey imaging system (LI-COR Biosciences). Membranes were stripped between blots by incubation in stripping buffer (Thermo Fisher) for 15 min and were then washed three times for 5□min each time with PBS.

### Luciferase Assays

HeLa cells were transiently transfected, as described above. PKR or MLKL promoter conjugated luciferase reporter constructs (Table S4) were used in combination with either TgNSM-Ty expression or empty vector as a control. The Renilla reporter plasmid pRL-TK (Promega) was co-transfected as a control for transfection efficiency. Twenty-four hours later, transfected cells were treated with human IFN-γ at (100 U/mL for 24 hr) or human IFN-β (1000 U/ml) and firefly luciferase activity was determined using the dual luciferase reporter assay system (Promega) on Cytation 3 (BioTek) multimode plate imager according to the manufacturer’s protocol. Experiments were repeated at least three times.

For GAS and ISRE interferon response in HeLa cells, 5×-GAS-Gaussia luciferase or 11 ×-ISRE-Gaussia luciferase constructs (Table S4) were used in combination with either TgNSM-Ty expression or empty vector as a control. Twenty-four hours after transfection, cells were treated with human IFN-γ at (100 U/mL for 24 hr) or human IFN-β (1000 U/ml) and gaussia luciferase activity was determined using Gaussia Luciferase Glow Assay Kit (Thermo) on Cytation 3 (BioTek) multimode plate imager according to the manufacturer’s protocol. Experiments were repeated at least three times.

### Flow Cytometry and RNA purification

Bradyzoites were differentiated in HFF cells cultured in T25 flasks. The monolayers were infected with 0.2×10^6^ parasites expressing mNeon under the control of BAG1 promoter and mCardinal under TUB promoter per T25 flask in D10 medium for 2 hr at 37°C to allow for invasion. For high pH bradyzoite induction, the media was changed to RPMI-HEPES (Thermo Fisher) lacking sodium bicarbonate pH 8.2. then grown for an additional 5 days at 37°C at ambient CO_2_ levels (~0.03% CO_2_). The media was changed every 2 days (to prevent acidification). Bradyzoites were stimulated with human IFN-γ (R&D systems) at final experimental concentration of 100 U/ml for 6 hr after 5 days in differentiation media prior to being washed with PBS and dislodged with trypsin. A minimum of 3×10^3^ mNeon/mCard positive HFF cells were collected into RT lysis buffer + 10 μl/ml 2-mercaptoethanol using a Sony SY3200 “Synergy cell sorter (Sony). RNA was extracted and purified from all samples using the RNeasy Mini kit (QIAGEN).

### Library Preparation and Sequencing

Total RNA from three independent biological replicates was submitted to the Genome Access Technology Center (Washington University School of Medicine) for library prep and sequencing. Total RNA integrity was determined using Agilent Bioanalyzer. Library preparation was performed with 1-10ng of total RNA. ds-cDNA was prepared using the SMARTer Ultra Low RNA kit for Illumina Sequencing (Takara-Clontech) per manufacturer’s protocol. cDNA was fragmented using a Covaris E220 sonicator using peak incident power 18, duty factor 20%, cycles per burst 50 for 120 seconds. cDNA was blunt ended, had an A base added to the 3’ ends, and then had Illumina sequencing adapters ligated to the ends. Ligated fragments were then amplified for 15 cycles using primers incorporating unique dual index tags. Fragments were sequenced on an Illumina NovaSeq-6000 using paired end reads extending 150 bases.

### Analysis of RNA-Seq Data

Demultiplexed fastq files were imported into Partek Flow (Partek, Inc.) Reads were then mapped to the Homosapiens hg38 genome build (NCBI GenBank assembly ID GCA_000001405.15) using the STAR aligner with default parameters (Dobin et al., 2013), and the number of reads per gene was quantified based on the human Ensembl Transcripts release 100. For differential expression analyses, we first normalized gene expression values by dividing the number of reads per gene by the total number of reads per sample to obtain counts per million (CPM) values. Pair-wise comparisons of sample types (3 biological replicates each) were performed using the Partek genespecific analysis (GSA) algorithm, which is a multimodel approach that determines the best response distribution (e.g. lognormal, Poisson, or negative binomial) for each gene based on Akaike information criteria (AICc) model selection rather than fitting all data to one distribution. Genes were not included in the analysis if their mean CPM value over all samples was less than 1. Genes were considered significantly differentially expressed between WT and the deletion strains (Δ*nsm*, Δ*ist*, Δ*nsm*/Δ*ist*) if the P value was less than 0.05 and the absolute fold change was greater than 1.5 and used for downstream for Reactome pathway analysis (Jassal et al., 2020).

### Real-Time PCR

Samples were prepared as described above in section Flow Cytometry and RNA purification. Complementary DNA (cDNA) was prepared using a Bio-Rad iScript cDNA Synthesis Kit (Bio-Rad Laboratories, Inc.) as per the manufacturer’s instructions. Realtime PCR was performed using TB Green^®^ Advantage^®^ qPCR premix (Takara Bio USA, Inc.) as per the manufacturer’s instructions. Data acquisition was done in QuantStudio3 (Applied Biosystems) and analyzed using QuantStudio Design and Analysis Software (Applied Biosystems). Primers are listed in (Table S5). Comparative cycle threshold values were used to evaluate fold change in transcripts using the β-Actin gene as an internal transcript control.

## QUANTIFICATION AND STATISTICAL ANALYSIS

All data were collected and analyzed without blinding. Data was analyzed using Prism software (version 7.01; Graphpad). Parametric statistical tests were used when the data followed an approximately Gaussian distribution, non-parametric tests were used when populations were clearly not Gaussian. Comparisons were considered statistically significant when *P* values were less than 0.05. Experiment-specific statistical information is provided in the figure legends or associated method details including replicates (n), trials (N), standard error, and statistical test performed.

## Supplemental items

**Table S1** Summary of Mass Spectrometry Analysis of APEX2 experiments. Related to Figure 1

**Table S2** Summary of Mass Spectrometry Analysis of Human and Toxoplasma Proteins Immunoprecipitated with TgNSM-Ty. Related to Figure 3.

**Table S5** Oligonucleotides Used in This Study. Related to STAR Methods.

