## Supplemental materials for "*Toxoplasma gondii* secreted effectors co-opt host repressor complexes to inhibit necroptosis"

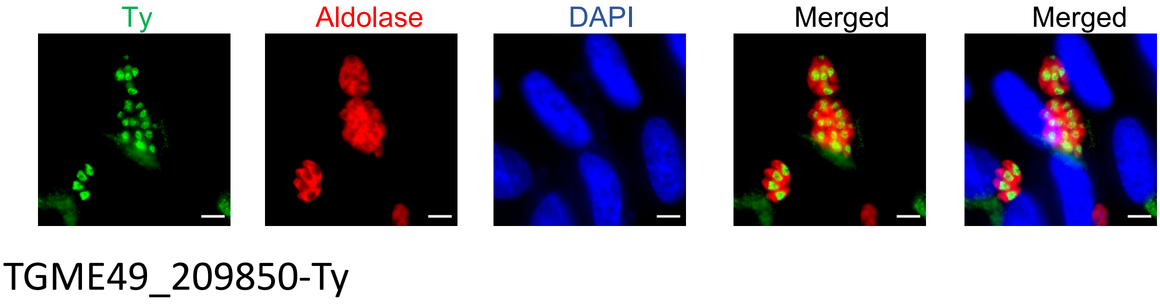


**Figure S1 TGME49_209850 localizes to the *T. gondii* nucleus. Related to Figure 1.**

HFF cells infected with type I (RH) endogenously tagged parasites TGME49_209850-Ty. Cells were fixed 24 hr post-infection, stained with mouse anti-Ty and anti-mouse IgG Alexa Fluor 488 (green), rabbit anti-Aldolase and rabbit IgG Alexa Fluor 568 (red) to detect all parasites, and DAPI (blue). Scale bars = 5 µm

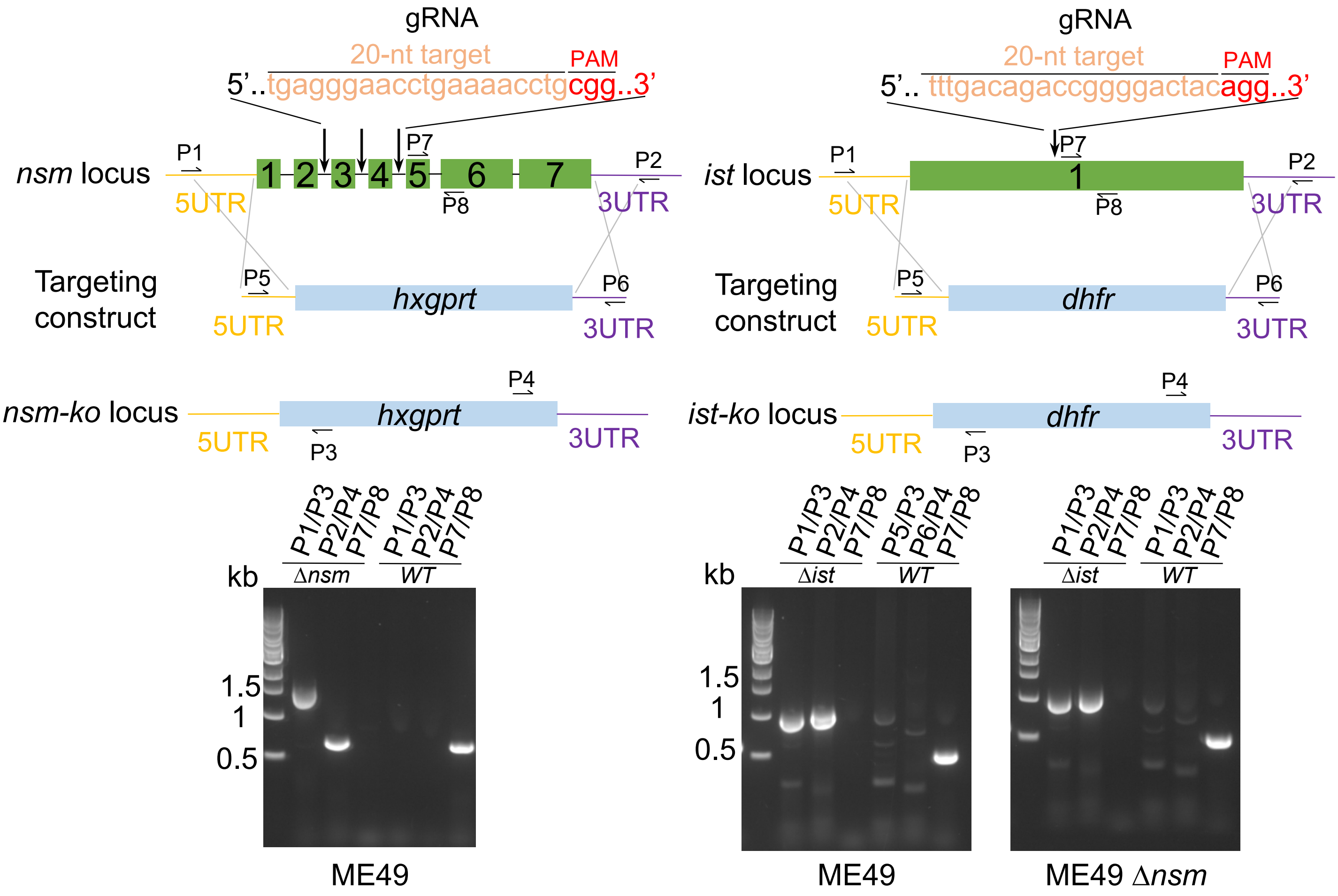


**Figure S2 Strategy for generation of knockout lines. Related to Figure 4**

Schematic representation of the strategy for CRISPR/Cas9-mediated gene deletion used to generate transgenic strains used in this study. A single sgRNA expressing CRISPR/Cas9 (gRNA) plasmids targeting the middle of the genes were used to mediate a knockout by homologous recombination. Targeting constructs consisted of a selection cassette (Δnsm-hxgprt, Δist-dhfr) and long homology flanks (~500bp) immediately upstream of the translation initiation site (left arm) and downstream of the stop codon (right arm) as homologous arms to the flanking regions of the GOI. Diagnostic PCRs results (bottom panel) to verify locus disruption. The priming sites for PCR primers are indicated: P1/P3 and P2/4 confirm integration of left and right homologous arms, respectively; P7/8 examines the integrity of the endogenous gene. A successful knockout clone gave positive PCR products in P1/P3 and P2/4 but no product in P7/8, whereas the wild type parasites gave the opposite. See also Table S5 for oligonucleotide sequences.

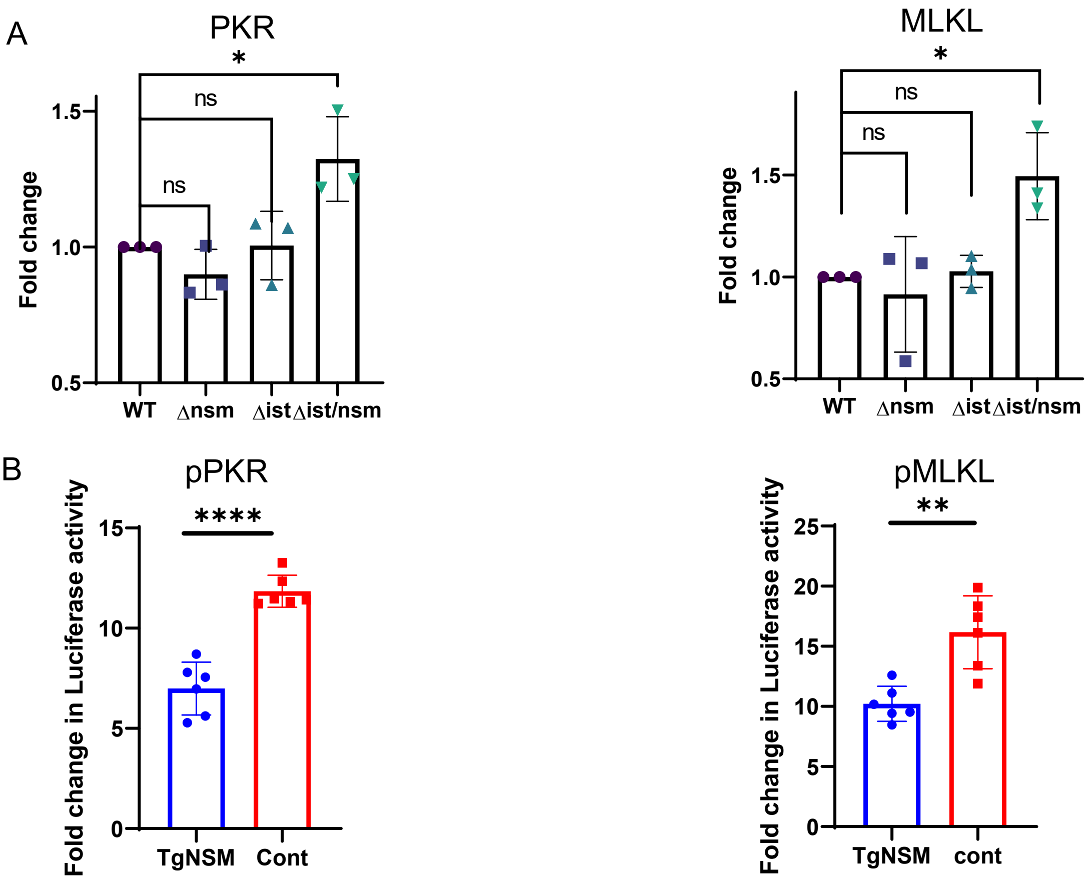


**Figure S3 TgNSM together with TgIST block transcription of PKR and MLKL upon IFN-β stimulation. Related to Figure 4**

(A) qPCR showing fold induction of mRNA transcripts in HFFs infected with parasite strains carrying pTUB-mCardinal and pBAG1-mNeonGreen reporters, grown in HFF cells under alkaline stress for 5 days, followed by treatment with IFN-β (1000 U/mL for 6 hr) purified by FACS based on the presence of both reporters. Comparative cycle threshold (Ct) values were used to evaluate the fold change in transcripts using b-actin (ActB) as an internal transcript control. Data are plotted as fold change ± SEM from at least 3 independent experiments per gene. There were significant differences between the compared groups *$P < 0.05$ using one-way ANOVA with Tukey's multiple comparison test.

(B) MLKL and PKR luciferase promotor constructs were transiently transfected into HeLa cells with TgNSM-Ty or empty vector. Twenty-four hr later, transfected cells were treated with IFN-β at (1,000 U/mL for 24 hr) and firefly luciferase activity was determined. The transfection efficiency was normalized against the-Rennila luciferase activity from the cotransfected pRL-TK vector. Results shown are fold induction over vector control and represent the averages and standard deviation from 3 biological replicates. Mean ± SD (n = 3 experiments, each with 3 replicates). **$P < 0.01$ and **** $P < 0.0001$, two-sample unpaired Student's t test.

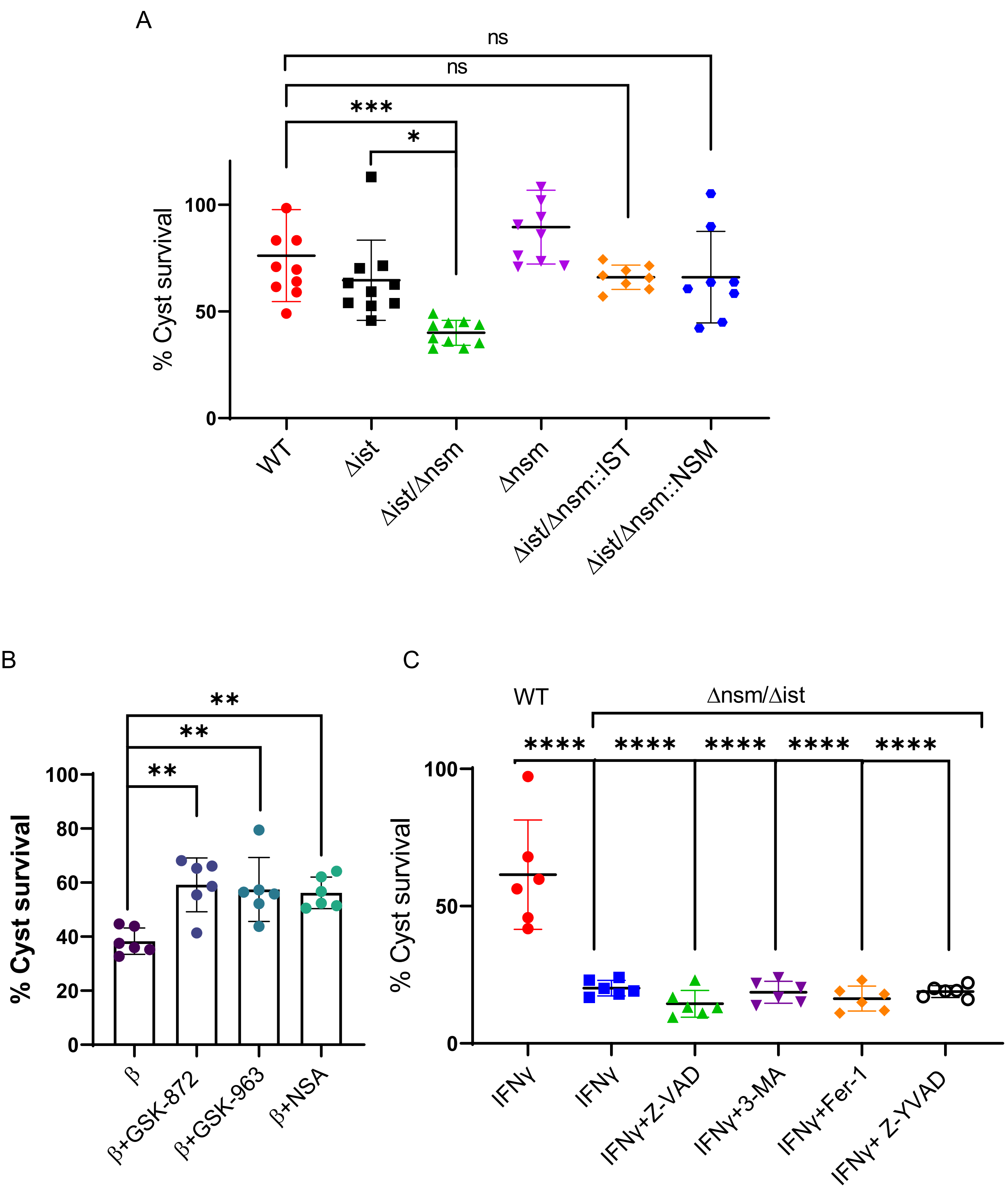
A
ns
ns
***
*
% Cyst survival
100
50
0
WT
Δist
Δist/Δnsm
Δnsm
Δist/Δnsm::IST
Δist/Δnsm::NSM
B
**
**
**
% Cyst survival
100
80
60
40
20
0
β
β+GSK-872
β+GSK-963
β+NSA
C
WT
Δnsm/Δist
****
****
****
****
****
% Cyst survival
100
50
0
IFNγ
IFNγ
IFNγ+Z-VAD
IFNγ+3-MA
IFNγ+Fer-1
IFNγ+ Z-YVAD

**Figure S4 Deletion of TgNSM and TgIST induces necroptotic host cell death and cyst rupture upon IFN-β stimulation. Related to Figure 5**

(A) Parasites were differentiated under alkaline conditions for 5 days before IFN-β 1000 U/ml treatment in presence of PI 50 µg/ml or PI only in control, and subsequent fixation and staining. Results shown are survival of cysts upon IFN-β treatment, calculated as percent of intact/PI negative cysts compared to PI only treated control and represent the averages and standard deviation from at least 3 biological replicates with each dot representing a technical duplicate, Mean ± SD (n = 3 experiments, each with 2 replicates 50-100 parasite cysts were counted in each treatment). *$P < 0.05$, *** $P < 0.001$, one-way ANOVA test with Tukey's multiple comparison test.

(B) ME49 Wild type and Δ*nsm/*Δ*ist* parasites were differentiated under alkaline conditions for 5 days before IFN-γ 100 U/ml treatment in presence of PI 50 µg/ml and Z-VAD 50 µM, Fer-1 0.5 µM, 3-MA 5mM, Z-YVAD 10 µM. PI and DMSO 0.05% only were used in control wells. Results shown are, survival of cysts upon IFN-γ treatment, calculated as percent of intact/PI negative cysts compared to PI only treated control and represent the averages and standard deviation from at least 3 biological replicates with each dot representing a technical duplicate Mean ± SD (n = 3 experiments, each with 2 replicates 50-100 parasite cysts were counted in each treatment). **** $P < 0.0001$, one-way ANOVA test with Dunnett's multiple comparison test.

(C) ME49 Δ*nsm/*Δ*ist* parasites were differentiated under alkaline conditions for 5 days before IFN-β 1,000 U/ml treatment in presence PI 50 µg/ml and RIPK1 inhibitor GSK-963 1 µM GSK-872 5 µM or NSA 10 µM. PI and DMSO 0.05% were used in control wells. Results shown are, survival of cysts upon IFN-β treatment, calculated as percent of intact/PI negative cysts compared to PI only treated control and represent the averages and standard deviation from at least 3 biological replicates with each dot representing a technical duplicate Mean ± SD (n = 3 experiments, each with 2 replicates 50-100 parasite cysts were counted in each treatment). **$P < 0.01$, one-way ANOVA test with Dunnett's multiple comparison test.

**Table S3 – *Toxoplasma gondii* lines used in this study. Related to STAR Methods**

| Line | Genotype | Source |
|---|---|---|
| RH (WT) | RH*Δhxgprt**Δku80* | (Huynh and Carruthers, 2009) |
| RH TgNSM-Ty | RH*Δhxgprt**Δku80*; TgNSM-Ty, DHFR-TS | This study |
| RH HCE1/TEEGR-Ty | RH*Δhxgprt**Δku80*; HCE1/TEEGR-Ty, DHFR-TS | This study |
| RH TGME49_209850 -Ty | RH*Δhxgprt**Δku80*; TGME49_209850 -Ty, DHFR-TS | This study |
| ME49 (WT) | ME49 *Δhxpgrt::FLUC* | (Tobin and Knoll, 2012) |
| ME49 *Δku80* | ME49 *Δhxpgrt::FLUC; Δku80::SAG1:CAT* | (Brown and Sibley, 2018) |
| ME49 TgNSM-Ty | ME49 Δhxpgrt::FLUC *Δku80::SAG1:CAT*; TgNSM-Ty, DHFR-TS | This study |
| ME49 *Δnsm* | ME49 Δhxpgrt::FLUC *Δnsm:: DHFR-TS*:*HXGPRT* | This study |
| ME49 *Δist* | ME49 Δhxpgrt::FLUC; *Δist::DHFR-TS* | This study |
| ME49 *Δnsm, Δist* | ME49 Δhxpgrt::FLUC; *Δnsm:: DHFR-TS*:*HXGPRT, Δist::DHFR-TS* | This study |
| ME49 TUB1:mCardinal BAG1:mNeonGreen | ME49 Δhxpgrt::FLUC; *TUB1:mCardinal, BAG1:mNeonGreen, SAG1:CAT* | This study |
| ME49 *Δnsm,* TUB1:mCardinal BAG1:mNeonGreen | ME49 Δhxpgrt::FLUC; *Δnsm:: HXGPRT; TUB1:mCardinal, BAG1:mNeonGreen, SAG1:CAT* | This study |
| ME49 *Δist,* TUB1:mCardinal BAG1:mNeonGreen | ME49 Δhxpgrt::FLUC; *Δist::DHFR-TS; TUB1:mCardinal, BAG1:mNeonGreen, SAG1:CAT* | This study |
| ME49 *Δnsm, Δist,* TUB1:mCardinal BAG1:mNeonGreen | ME49 Δhxpgrt::FLUC; *Δnsm:: HXGPRT; Δist::DHFR-TS; TUB1:mCardinal, BAG1:mNeonGreen, SAG1:CAT* | This study |
| ME49 *Δnsm, Δist/TgNSM-Ty* | ME49 Δhxpgrt::FLUC; *Δnsm:: DHFR-TS*:*HXGPRT, Δist::DHFR-TS,TgNSM-Ty CAT* | This study |
| ME49 *Δnsm, Δist/TgIST-Ty* | ME49 Δhxpgrt::FLUC; *Δnsm:: DHFR-TS*:*HXGPRT, Δist::DHFR-TS,TgIST-Ty CAT* | This study |

**Table S4 Plasmids used in this study, related to STAR Methods.**

| Common Name | Description | Source |
|---|---|---|
| pSAG1:CAS9-GFP, U6:sgUPRT | Template for making GOI targeting Cas9 plasmids | Addgene (#54467) (Shen et al., 2014) |
| pSAG1:CAS9-GFP, U6:sgHCE1/TEEGR (3') | CRISPR plasmid targeting HCE1/TEEGR 3' UTR for C-terminal tagging | This study |
| pSAG1:CAS9-GFP, U6:sg209850 (3') | CRISPR plasmid targeting ME49_209850 3' UTR for C-terminal tagging | This study |
| pSAG1:CAS9-GFP, U6:sgTgNSM (3') | CRISPR plasmid targeting TgNSM 3' UTR for C-terminal tagging | This study |
| pSAG1:CAS9-GFP, U6:sgTgNSM | CRISPR plasmid targeting TgNSM coding sequence for gene deletion | This study |
| pSAG1:CAS9-GFP, U6:sgDHFR | CRISPR plasmid targeting DHFR resistance cassette coding sequence for TgIST complementation | This study |
| pSAG1:CAS9-GFP, U6:sgHX | CRISPR plasmid targeting HX resistance cassette coding sequence for TgNSM complementation | This study |
| pUC19 | Subcloning; for generating *TgNSM-Ty* and TgIST-Ty complement plasmids. | New England Biolabs, Inc. Cat# N3041S |
| pFloxed DHFR-TS* | Template for amplification of DHFR resistance | In-house |

| | | |
|---|---|---|
| | cassette for obtaining deletion strains | |
| pLinker-2xTy-HXGPRT-LoxP | Template for amplification of HX resistance cassette for obtaining deletion strains and for obtaining endogenously TY tagged proteins | In-house |
| pLinker-2xTy-DHFR-LoxP | Template for amplification of DHFR resistance cassette for obtaining endogenously TY tagged proteins | In-house |
| pTUB1-mCardinal-pSag1-CAT | Subcloning; for generating pTUB1-mCardinal-pSag1-CAT-pBAGmNeonGreen plasmid. | In-house |
| pTubLinker-TYx3-mNeonGreen-TYx3-HX(floxed) | Template for amplification of mNeonGreen for obtaining pTUB1-mCardinal-pSag-CAT-pBAGmNeonGreen plasmid | In-house |
| pNJ_26-pBAG1-mCherry | Template for amplification of BAG1 5’ UTR for obtaining pTUB1-mCardinal-pSag-CAT-pBAGmNeonGreen plasmid | In-house |
| pBS-SAG1-CAT | Template for amplification of CAT resistance cassette for obtaining pTgNSM-Ty-complement-CAT and pTgIST-Ty-complement-CAT plasmids | In-house |

| | | |
|---|---|---|
| pTUB1-mCardinal-pSag1-CAT-pBAGmNeonGreen | Reporter for pTUB1-mCradinal and pBAG1-mNeonGreen expression | This study |
| pUPRT-5UTR-TgIST-TY-3UTR-DHFR | Template for amplification of TgIST-TY for obtaining pTgIST-Ty-complement-CAT plamid | (Olias et al., 2016) |
| pTgNSM-Ty-complement-CAT | *TgNSM-2Ty* fusion with *CAT* drug selectable marker flanked by homology arms from *TgNSM*. Used with pSAG1:CAS9-GFP, U6:sgHX to obtain TgNSM complement strain | This study |
| pTgIST-Ty-complement-CAT | *TgIST-2Ty* fusion with *CAT* drug selectable marker flanked by homology arms from *TgIST*. Used with pSAG1:CAS9-GFP, U6:sgDHFR to obtain TgIST complement strain | This study |
| pSG5HA-mSMRTα (a.a. 1-2470) | Full length HA tagged SMRT protein expression vector | (Varlakhanova et al., 2011) Gift from Dr. Martin L. Privalsky |
| pSG5HA-mNCoR (a.a. 1-2453) | Full length HA tagged NCoR protein expression vector | Gift from Dr. Martin L. Privalsky |
| pN1-TBLR1-mCherry | Full length TBLR1 fused to mCherry protein expression vector | (Kruusvee et al., 2017) |
| pN1-TBL1-mCherry | Full length TBL1 fused to mCherry protein expression vector | (Kruusvee et al., 2017) |
| pcDNA3-H2B-V5-APEX2 | Nuclear APEX2 fused to H2B protein | (Lee et al., 2016) |

| | | |
|---|---|---|
| pcDNA3 APEX2-NES | Cytoplasmic APEX2 | Addgene #49386 (Lam et al., 2015) |
| pSBbi-GP | SB-transposon with constitutive bi-directional promoter | Addgene #60511 (Kowarz et al., 2015) |
| pSBbi-GP-TgNSM-TY | TgNSM fused to TY expression vector | This paper |
| pGL3-pPKR-luciferase | PKR promoter driving Firefly luciferase expression | (Yoon et al., 2009) |
| pGL4-pMLKL-luciferase | MLKL promoter driving Firefly luciferase expression | (Gunther et al., 2016) |
| 5×-GAS-Gaussia luciferase | GAS response element driving Gaussia luciferase | (Matta et al., 2019) |
| 11×-ISRE-Gaussia luciferase | ISRE response element driving Gaussia luciferase | (Matta et al., 2019) |
| pRL-TK | Mammalian co-reporter vector for the weak constitutive expression of wild-type Renilla luciferase | Promega cat# E2231 |
